# Identification of a Nonsense-Mediated Decay pathway at the Endoplasmic Reticulum

**DOI:** 10.1101/2020.02.24.954453

**Authors:** Dasa Longman, Kathryn A. Jackson-Jones, Magdalena M. Maslon, Laura C. Murphy, Robert S. Young, Jack J. Stoddart, Martin S. Taylor, Dimitrios K. Papadopoulos, Javier F. Cáceres

**Affiliations:** MRC Human Genetics Unit, Institute of Genetics and Molecular Medicine, University of Edinburgh, Crewe Road South, Edinburgh EH4 2XU, UK; Centre for Global Health Research, Usher Institute, University of Edinburgh, Old Medical School, Teviot Place, Edinburgh EH8 9AG, UK

**Keywords:** Nonsense-mediated decay (NMD), RNA quality control, UPF1, NBAS, Syntaxin 18, ER stress, UPR

## Abstract

Nonsense-mediated decay (NMD) is a translation-dependent RNA quality control mechanism that occurs in the cytoplasm. However, it is unknown how NMD regulates the stability of RNAs translated at the Endoplasmic Reticulum (ER). Here, we identify a localized NMD pathway dedicated to ER-translated mRNAs. We previously identified NBAS, a component of the Syntaxin 18 complex involved in Golgi-to-ER trafficking, as a novel NMD factor. Here, we show that NBAS fulfils an independent function in NMD. This ER-NMD pathway requires the interaction of NBAS with the core NMD factor UPF1, which is partially localized at the ER in the proximity of the translocon. NBAS and UPF1 co-regulate the stability of ER-associated transcripts, in particular those associated with the cellular stress response. We propose a model where NBAS recruits UPF1 to the membrane of the ER and activates an ER-dedicated NMD pathway, thus providing an ER protective function by ensuring quality control of ER-translated mRNAs.

**Graphical Abstract:** 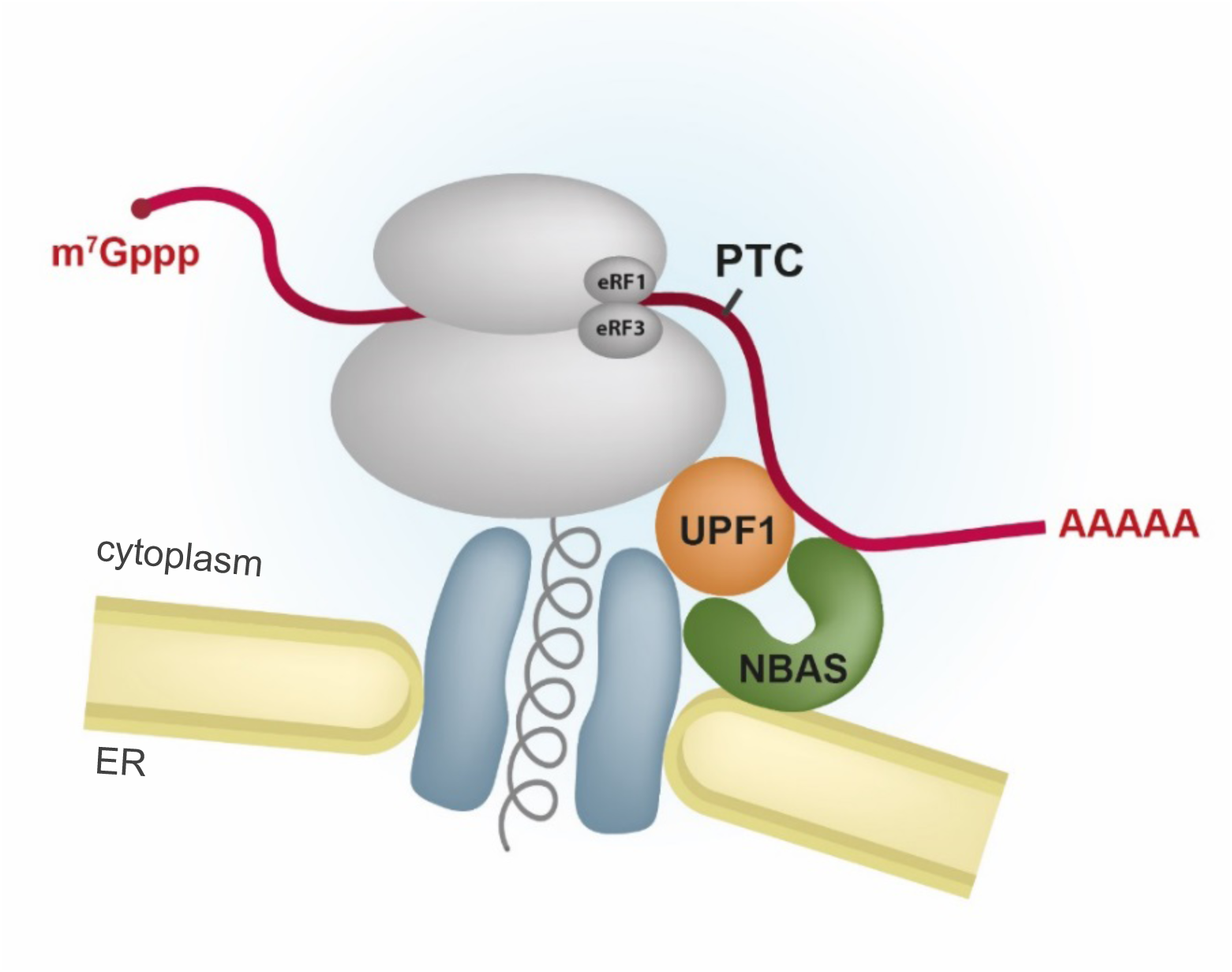

**HIGHLIGHTS:**

- NBAS is an NMD factor that localizes to the membrane of the endoplasmic reticulum (ER)
- NBAS has dual, independent, roles in Golgi-to-ER retrograde transport and in ER-NMD
- NBAS recruits the core NMD factor UPF1 to the membrane of the ER
- The ER-NMD pathway targets for degradation mRNAs that are translated at the ER

## Introduction

The nonsense-mediated decay (NMD) pathway is a highly conserved surveillance mechanism that targets mRNAs harboring premature termination codons (PTCs) for degradation. In doing so, it prevents the accumulation of truncated proteins and modulates the phenotypic outcome of genetic disorders that arise due to the presence of PTCs (Bhuvanagiri et al., 2010). Importantly, NMD also controls the stability of a large number of endogenous RNAs and fine-tunes many physiological processes (reviewed by (Karousis and Mühlemann, 2019; Kurosaki et al., 2019)).

Core NMD factors were initially identified in genetic screens in *C. elegans* and in *S. cerevisiae* and were later shown to be involved in NMD in other species, including *Arabidopsis*, *Drosophila* and mammals. Human orthologs of these core NMD factors are SMG1, UPF1, UPF2, UPF3, SMG5, SMG6, and SMG7 (Nicholson et al., 2010). Additional NMD factors have been identified using a variety of different experimental approaches, including interactome studies, RNAi screens in nematodes and CRISPR screens in mammalian cells (Alexandrov et al., 2017; Baird et al., 2018; Hug et al., 2016). Mechanistically, NMD is tightly linked to mRNA translation and is initiated by the recognition of a PTC by the surveillance (SURF) complex within which the RNA helicase UPF1, and its associated kinase, SMG1, bind to the ribosomal release factors eRF1 and eRF3 (He and Jacobson, 2015; Kishor et al., 2019). Subsequently, components of the SURF complex interact with the core NMD factors UPF2 and UPF3b, and with an exon junction complex (EJC) located downstream of the PTC, to form the decay-inducing complex (DECID) that triggers UPF1 phosphorylation by SMG1, and dissociation of eRF1 and eRF3 (Buchwald et al., 2010; Kashima et al., 2006; Singh et al., 2007). This leads to the recruitment of mRNA degradation factors that trigger RNA decay. Substrate selection for NMD occurs not only in an EJC-dependent manner, but alternatively via an EJC-independent mechanism, which targets transcripts harboring very long 3’UTRs (Kurosaki et al., 2019; Metze et al., 2013).

Using RNAi screens in *C. elegans*, we previously identified several novel NMD factors that were also essential for viability, suggesting that they fulfil other cellular functions in nematodes, where this pathway is not essential (Casadio et al., 2015; Longman et al., 2007). One of these novel NMD factors is encoded by *smgl-1* that corresponds to the human gene *NBAS* (neuroblastoma amplified sequence, also known as *NAG,* for neuroblastoma amplified gene). *NBAS* was first identified as a gene that is co-amplified with the *N-myc* gene in human neuroblastomas, however no clear role in the disease has been reported (Scott et al., 2003; Wimmer et al., 1999). We went on to show that NBAS acts in concert with UPF1 to co-regulate a large number of transcripts not only in nematodes but also in zebrafish and human cells (Anastasaki et al., 2011; Longman et al., 2013). Of interest, *NBAS* encodes a peripheral ER membrane protein that is a component of the Syntaxin 18 complex, which functions in Golgi-to-ER retrograde transport (Aoki et al., 2009). Recently, a series of loss-of-function mutations in *NBAS* have been found in several human conditions, including biallelic mutations in *NBAS* in patients with a multisystem disease involving liver, eye, immune system, connective tissue and bone (Haack et al., 2015; Segarra et al., 2015). Compound heterozygous variants in *NBAS* were also identified as a cause of atypical osteogenesis imperfecta (Balasubramanian et al., 2017) and in a short stature with optic atrophy and Pelger-Huët anomaly (SOPH) syndrome (Maksimova et al., 2010). Currently, it remains unclear whether the phenotypes observed in patients with mutations in *NBAS*, are due to a compromised NMD response, defects in Golgi-to-ER retrograde transport, or a combination of both.

Despite initial controversy concerning the intracellular location of NMD in mammalian cells, it has been conclusively demonstrated that decay of a PTC-containing β-globin NMD reporter occurs in the cytoplasm (Trcek et al., 2013). The ER is a major site of localized protein synthesis, with approximately a third of all mRNAs being translated there, in particular, those encoding proteins entering the secretory pathway. It has become increasingly evident that ER and cytosol constitute different environment for protein translation and post-transcriptional gene regulation (Reid and Nicchitta, 2015). Current efforts have focused on the mechanism and regulation of cytoplasmic NMD; however, it is largely unknown how this mechanism operates on mRNAs that are translated at the ER, which due to their intrinsic localized translation will not have sufficient exposure to cytoplasmic NMD. There is a precedent for a localized NMD response, as seen in neurons, where NMD regulates the expression of both dendritic and axonal mRNAs upon their activation of localized mRNA translation (Colak et al., 2013; Giorgi et al., 2007). Both NBAS and a second novel NMD factor identified in our RNAi screens, SEC13 (nuclear pore and COPII coat complex component), localize to the membrane of the ER, raising the possibility that they could be involved in an ER-localized NMD pathway (Casadio et al., 2015; Longman et al., 2007).

Here, we present evidence that reveals a central role for NBAS, acting together with UPF1, in an NMD response that is associated with the ER. We show that NBAS has dual roles in NMD and Golgi-to-ER retrograde transport; but importantly, these functions act independent of each other. We demonstrate that NBAS recruits the core NMD factor UPF1 to the membrane of the ER and promotes the degradation of NMD substrates that are translated at the ER.

## RESULTS

### A dual role of NBAS in Golgi-to-ER transport and NMD

*NBAS* encodes a 2,371 amino acid protein containing WD40 repeats and a SEC39 domain, present in proteins involved in the secretory pathway (Figure 1A). Together with RINT1 and ZW10, NBAS forms part of the evolutionarily conserved NRZ complex, which functions as a tethering complex for retrograde trafficking of COPI vesicles from the Golgi to the ER and is part of the larger Syntaxin 18 complex (Aoki et al., 2009; Civril et al., 2010).

**Figure 1.**
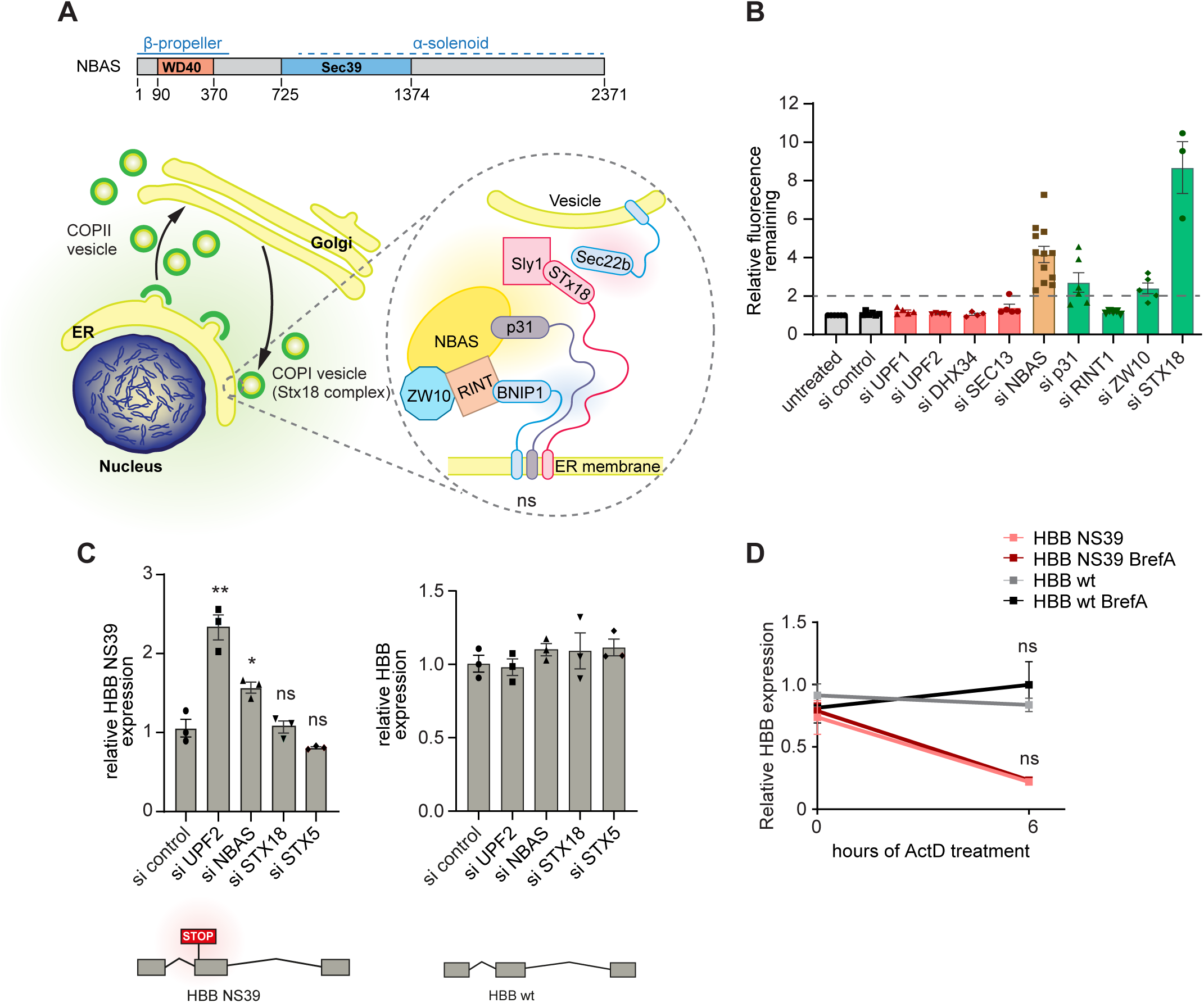
NBAS is an ER-localized protein with dual, independent, functions in NMD and ER secretion. (A) Cartoon depicting the functional domains of NBAS and its intracellular localization at the Endoplasmic Reticulum (ER), as part of the Syntaxin 18 complex. (B) NMD activity is not required for constitutive secretion in HeLa C1 cells. Depletion of NMD factors, with the exception of NBAS, does not affect the ability of C1 cells to secrete a GFP-based reporter construct. GFP accumulation was measured by flow cytometry and a secretion defect was defined as a two-fold accumulation of GFP reporter (after the addition of 1 µM D/D Solubilizer (ligand) compared to untreated C1 cells (dashed line)). Each point represents one biological replica, bars indicate mean with SEM. See Figure S1A for more details. (C) NMD activity is not affected by depletion of STX18 and STX5, factors that are required for constitutive secretion. In contrast, depletion of UPF2 and NBAS led to an upregulation of the HBB NS39 NMD-sensitive reporter. The expression of the control HBB wt reporter that is not an NMD substrate was not changed. HeLa cells stably expressing the HBB NS39 or wt reporters were depleted of the indicated factors and the steady state level of the reporter mRNA was measured by qRT-PCR and normalized to POLR2J expression. Each point represents one biological replica, bars indicate mean with SEM. Significance was determined by two-tailed unpaired t-test: **: p<0.005; *: p<0.05; ns: not significant. (D) NMD activity is not affected by blocking constitutive secretion. Treatment with Brefeldin A does not change the stability of HBB NS39 NMD-sensitive transcripts (compare red and pink lines). HeLa cells carrying HBB wt or HBB NS39 transcripts were treated with Actinomycin D to block transcription, and Brefeldin A to block constitutive secretion. HBB expression was measured as described in panel (C). Each point represents the mean and SEM of three biological replicas. The significance of Brefeldin A treatment was determined by two-tailed unpaired t-test: ns: not significant. See Figure S1C for the effect of Brefeldin A on secretion.

To dissect whether NBAS had separate roles in NMD and Golgi-to-ER transport, or whether these processes influence each other, we first tested whether abrogating the NMD pathway influences ER trafficking. We used a modified flow cytometry-based assay that relies on the expression of an eGFP fluorescent reporter for measuring constitutive secretion (Gordon et al., 2010). This assay is based on the property of mutant FKBP proteins (F36M) to form large aggregates that when expressed in the ER cannot be secreted. However, upon incubation with an FKBP (F36M) ligand, D/D Solubilizer, these aggregates are solubilized leading to efficient secretion (Gordon et al., 2010). We used this reporter in combination with siRNA-mediated knock-down of individual NMD factors or of components of the Syntaxin 18 complex (Figures 1B and S1A). Fluorescence of clonal HeLa C1 cells stably expressing the eGFP reporter decreased upon addition of the ligand, D/D Solubilizer, to the level of control cells, as was previously observed (Gordon et al., 2010) (Figure S1A, left panel). As expected, siRNA-mediated knockdown of components of the Syntaxin 18 complex, including STX18, p31 and NBAS resulted in an increase of the remaining fluorescence, due to an interference with the secretion process (Figure 1B). We observed some effect upon depletion of ZW10, but almost no effect with the knock-down of RINT1, perhaps reflecting a differential contribution of these components to the secretion process. Importantly, we observed that a strong reduction in the levels of individual NMD factors did not interfere with secretion. This was the case for UPF1 and UPF2, but also for the RNA helicase DHX34, and for SEC13, another NMD factor that localizes to the ER (Figures 1B and S1A). From this experiment, we conclude that NMD activity is not required for ER secretion.

Next, we investigated whether interfering with constitutive secretion had any effect on the NMD pathway. For this, HeLa cells stably expressing a well characterized β-globin NMD reporter harboring a nonsense mutation at position 39 (HBB NS39) (Trecartin et al., 1981), or its wild-type (wt) HBB counterpart, were depleted of NMD factors (UPF2 or NBAS) or of secretion factors (STX18 or STX5) using specific siRNAs (Figure S1D). As expected, the level of HBB WT mRNA remained unchanged upon UPF1, NBAS, STX18 or STX5 depletion (Figure 1C, right panel). By contrast, depletion of UPF2 or of NBAS resulted in an increased level of the HBB NS39 NMD reporter mRNA, compared to mock depleted cells (Figure 1C, left panel). Importantly, depletion of either STX18 or STX5 did not affect the level of β-globin NMD reporter mRNA (Figure 1C), indicating that normal ER secretion is not required for NMD. These results were confirmed using a fluorescent NMD reporter (NMD+) that quantifies NMD activity at the single cell level. Cells carrying the NMD+ reporter were identified by the constitutive expression of red fluorescence, whereas NMD activity was determined by the mean green fluorescence, which is subject to NMD, in all red cells (Pereverzev et al., 2015). Here again, whereas knock-down of UPF2 and NBAS resulted in increased levels of the NMD reporter (measured by increased green fluorescence), depletion of STX18 or STX5 had no effect (Figure S1B). We extended these observations to show that NMD activity is not affected by blocking constitutive secretion with the use of Brefeldin A that inhibits Golgi-to-ER protein transport (Aridor et al., 1995) (Figure S1C). HeLa cells stably expressing either WT HBB or HBB NS39 were treated with Actinomycin D to block transcription in the presence or absence of Brefeldin A. We observed that the stability of the HBB NS39 NMD reporter mRNA was not increased by blocking ER secretion (Figure 1D). Altogether, these experiments show that NMD activity and ER secretion are not functionally linked, strongly suggesting that NBAS has two independent roles in Golgi-ER retrograde transport and in NMD.

### NBAS regulates a subset of NMD targets specifically translated at the ER

Previously, we conducted RNA profiling experiments that revealed a large proportion of NBAS mRNA targets are co-regulated by the core NMD factor UPF1, with a significant enrichment for genes involved in the cellular stress response (Longman et al., 2013). Here, we extended this analysis by RNA-sequencing to profile changes in mRNA abundance upon depletion of NBAS or UPF1. Both NBAS and UPF1 were the most significantly downregulated genes in the relevant samples (fold change −1.85, p<2.3×10^-16^ and fold change −3.28, p<1.14×10^-148^ respectively)(Table S1). Depletion of UPF1 significantly affected the mRNAs of 4756 genes, whereas mRNAs of 2411 genes increased. Depletion of NBAS altered the mRNA levels of 421 genes (Table S1). Of these, 209 genes were increased in abundance, consistent with the view that NBAS regulates a subset of the UPF1 targets. We observed a robust co-regulation of mRNAs when UPF1 or NBAS was depleted (Pearson’s correlation r=0.67, p<2.2×10^-16^) (Figure 2A), indicating that UPF1 and NBAS function in a common pathway. Gene ontology (GO) analysis of the mRNAs regulated by both NBAS and UPF1 revealed a strong enrichment for the secretome, which is preferentially translated at the ER translocon (Table S2). Interestingly, we found a significant increase in fold change in genes associated with the ER unfolded protein response (UPR) (GO: 0006986) when either NBAS or UPF1 was depleted (p < 0.05 and p < 0.001 respectively; Wilcoxon rank sum test), showing that ER stress response is perturbed when ER-NMD is not functional. This supports our hypothesis that an ER-specific NMD pathway is essential for cellular homeostasis. To further characterize genes regulated by NBAS and UPF1, we intersected targets regulated by either NBAS or UPF1 and also common NBAS and UPF1 targets, with three experimental datasets identifying genes localized and/or translated at the ER. APEX-seq, a method for RNA sequencing based on direct proximity labeling of RNA led to the identification of 1077 mRNAs localized at the ER (Fazal et al., 2019), cell fractionation followed by ribosome footprinting coupled with deep-sequencing identified 486 mRNAs (Reid & Nicchitta, 2012), and proximity-specific ribosome profiling, based on ER membrane proximity labelling of ribosomes identified 686 mRNAs translated at the ER (Jan et al., 2014). We found that NBAS targets are strongly enriched for experimentally validated ER genes (31.6%), as compared to 14.4% of UPF1 targets. Furthermore, 18.2% of NBAS targets were found in more than two of the datasets, as compared to only 6.3% of UPF1 targets. Considering NBAS and UPF1 common targets, 37.9% of targets were found in at least one dataset, whilst 23.2% were found in more than two datasets (Figure 2B). NBAS-regulated targets were strongly enriched for experimentally identified ER-localized genes (OR=6.07, p<0.001, Fisher’s Exact test), whereas UPF1 targets were only modestly enriched (OR=2.44, p<0.001, Fisher’s Exact test). Compared to UPF1 targets, we found that those targets which were also regulated by NBAS were 2.6-fold more likely than expected by chance to be found at the ER (p<0.001, Exact Binomial test), confirming that NBAS and UPF1 together regulate NMD targets specifically at the ER.

**Figure 2.**
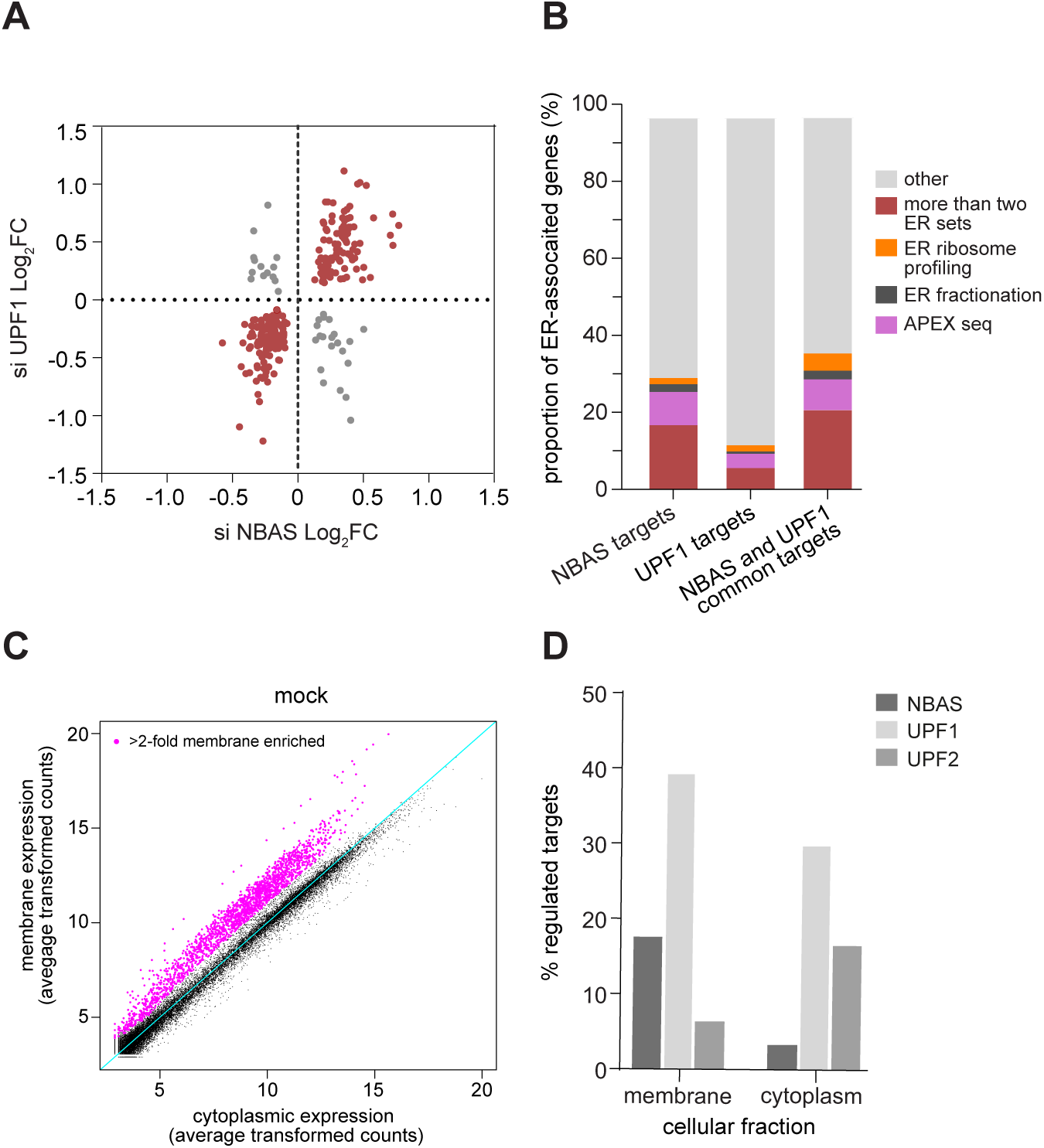
NBAS-regulated targets are enriched for ER-localized transcripts. (A) Scatter plot of differentially expressed transcripts shows a significant positive correlation between NBAS and UPF1-regulated targets ((Pearson’s correlation r= 0.67; p< 0.0001). (B) NBAS-regulated targets are enriched for experimentally identified ER-localized transcripts. Bar charts show the proportion of targets that are ER-localized by APEX-seq (Fazal et al., 2019), ER fractionation (Reid and Nicchitta, 2012) and ER proximity-specific ribosome profiling (Jan et al., 2014). The ER-association of NBAS targets is significantly higher from UPF1 targets (p< 0.0001, Binomial Exact test), whereas the difference between NBAS targets and NBAS-UPF1 common targets is not statistically significant (p= 0.1867, Binomial Exact test). (C) RNA-seq of samples from subcellular fractions (membrane and cytoplasm) allows identification of membrane-associated genes. Scatter plot shows average transformed counts of each gene in membrane and cytoplasmic fractions in mock-depleted cells. Differential expression analysis gave fold change values to expression between cytoplasm and membrane. Genes with 2-fold increased expression in the membrane fraction were assigned as “membrane-associated” (magenta points). All other genes were classified as “non-membrane” genes. Additional data referring to this panel can be found in Figure S2A. (D) NBAS preferentially regulates membrane-associated targets. Plot shows percentage of ‘membrane-associated’ genes in the membrane fraction and of ‘non-membrane’ genes in the cytoplasm that are regulated by each NMD factor. NBAS regulates over 5-fold higher percentage of membrane-associated genes than non-membrane genes (p<0.001, Exact Binomial test). By contrast, UPF1 regulates a similar percentage of both membrane-associated and non-membrane genes (1.3-fold higher percentage of membrane-associated genes, p<0.001, Exact Binomial test), and UPF2 regulates 2.5-fold higher percentage of non-membrane genes than membrane genes (p<0.001, Exact Binomial test). Genes were determined as regulated if they were significantly (p<0.05) increased in expression when the relevant factor was depleted. See Supplementary Figure 2B for the summary of regulated genes, and Supplementary Figure 2C for differential expression analysis.

We next examined how NBAS depletion affected transcripts specifically at the ER compared to the cytoplasm and how this differs from depletion of core NMD factors UPF1 and UPF2. We therefore performed RNA-sequencing on cytoplasmic and membrane fractions of HeLa cells depleted for each factor or mock-depleted control cells, following subcellular fractionation. By comparing expression of genes in each fraction in control samples we defined a group of “membrane-associated” genes that showed a 2-fold higher expression in the membrane fraction, whereas all other expressed genes were deemed “non-membrane” genes (Figure 2C). We assessed the validity of cell fractionation by identifying genes in the previously described experimentally validated ER datasets in our data. As expected, we observed that the majority of genes present in any two ER datasets were found in the membrane-associated set (OR = 66.94, p-value < 2.2e-16, Fishers Exact test) (Figure S2A). Next, we analyzed changes in gene expression of ‘membrane-associated’ genes in the membrane fraction, and of ‘non-membrane’ genes in the cytoplasmic fraction, upon depletion of individual NMD factors (Figures 2D, S2B and S2C). Knock-down of UPF1 led to a robust increase in the abundance of mRNAs in both membrane and cytoplasmic fractions, whereas depletion of NBAS led to an upregulation of a higher percentage of membrane-associated genes than of non-membrane genes (5-fold increase, p<0.001, Exact Binomial test). In contrast, depletion of UPF2 led to preferential upregulation of ‘non-membrane’ genes in the cytoplasm (> 2.5-fold, p<0.001, Exact Binomial test). Altogether, these results suggest that NBAS is a crucial component of an ER-NMD pathway that targets for degradation mRNAs that are translated at the ER, constituting a novel localized NMD response.

### Site of NMD for transcripts translated at the ER

Single-molecule RNA Fluorescent in Situ Hybridization (smRNA FISH) was previously used to localize the NMD response of a β-globin NMD reporter in single cells, leading to the conclusion that the NMD activity occurs in the cytoplasm (Trcek et al., 2013). Here, we have used single molecule RNA fluorescence in situ hybridization (smRNA FISH) to spatially map the location of NMD-mediated mRNA degradation of mRNAs that are translated at the ER and that we have also shown to be upregulated upon UPF1 and/or NBAS depletion (Longman et al., 2013). We used a set of fluorescent probes that label the full-length of two endogenous mRNAs targeted by NMD, which are translated either in the cytoplasm or at the ER. We selected the *SETD4* (SET Domain Containing 4) mRNA, which encodes a lysine methyl transferase and is translated in the cytoplasm (Faria et al., 2013). As an example of an mRNA that is translated at the ER we selected the *FAP* mRNA (also known as Seprase), which encodes Fibroblast Activation Protein alpha, a 170kDa membrane-bound gelatinase (Goldstein et al., 1997). Levels of *SETD4* mRNA were robustly increased upon knock-down of UPF1, whereas NBAS depletion had only a marginal stabilizing effect (Figure 3A). In contrast, we found that the ER-localized *FAP* mRNA was comparably upregulated upon knock-down of UPF1 and of NBAS (Figure 3A).

**Figure 3.**
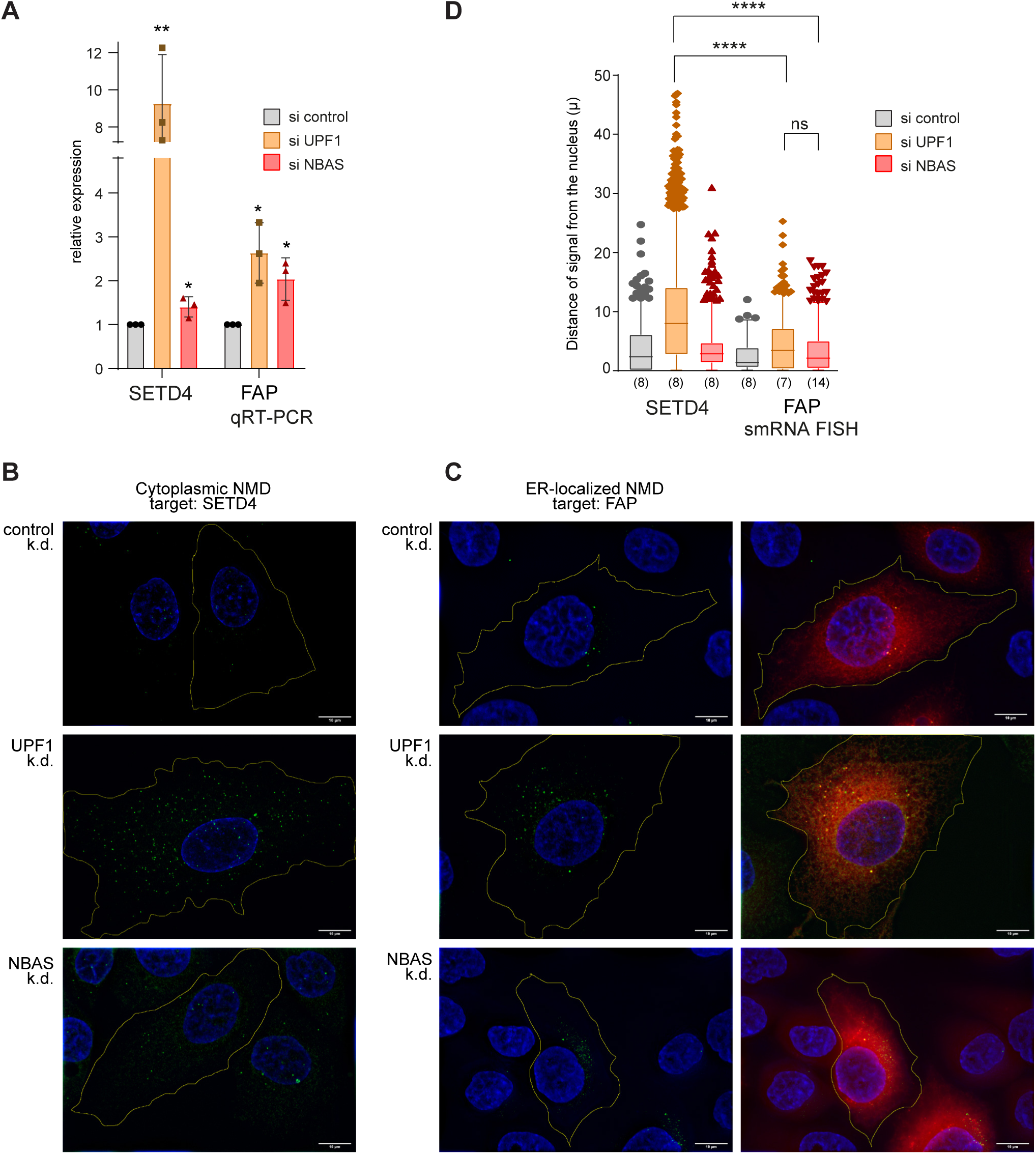
ER-localized NMD targets are degraded at the ER. (A) Relative changes in SETD4 and FAP expression upon UPF1 or NBAS depletion were measured by qRT-PCR and normalized to POLR2J expression. Each point represents one biological replica, bars indicate mean with SEM. Significance was determined by two-tailed unpaired t-test: **: p<0.005; *: p<0.05. (B-C) Single-molecule (sm) RNA FISH was used to visualize the site of accumulation of a cytoplasmic NMD target SETD4 (B), or an ER-localized NMD target FAP (C), within the cell when NMD was abrogated by either UPF1 or NBAS knock-down. Individual RNA molecules were visualized by a set of transcript-specific Quasar® 570-labelled Stellaris probes and pseudo-colored green. The cell outline was visualized by staining with MemBright 640nm probe and is indicated by the yellow line. The ER compartment was visualized by mEmeral-ER-3 co-transfected plasmid and is pseudocolored red (right column). Each scale bar is 10μm. (D) Distribution of FISH signal within imaged cells is represented by the distance of signal from the nucleus in each condition. Graph shows distances as 5-95 percentile box plots, numbers in brackets indicate the number of analyzed cells. Data was acquired in three independent experiments. Statistical significance was determined by one-way ANOVA test. ****: p_adj<0.0001; ns: not significant.

RNA FISH of *SETD4* mRNA in UPF1-depleted cells revealed a strong increase of uniformly distributed fluorescent signal throughout the cell, consistent with NMD taking place in the cytoplasm. As expected, depletion of NBAS led only to sporadic upregulation of SETD4 mRNA (Figure 3B). Importantly, RNA-FISH of the *FAP* mRNA in both UPF1 and NBAS-depleted cells revealed a strong increase of the fluorescent signal that clustered to the perinuclear region of the cell (Figure 3C, green signal). The RNA-FISH fluorescence overlapped with ER staining ((Figure 3C, right panels), thus spatially mapping the NMD response of an NBAS NMD target to the ER. In order to represent the differences in the distribution of the smRNA FISH signal within cells, we plotted the distance of the fluorescent FISH signal from the edge of the nucleus (defined by DAPI staining), for each experimental condition This demonstrated that UPF1 depletion leads to the accumulation of SETD4 mRNA that is distributed widely within the cell, which is significantly different from the distribution pattern of the ER-associated FAP mRNA signal that clusters close to the nuclear periphery upon depletion of both NBAS and UPF1 (Figure 3D). Altogether, these results provide strong evidence that NBAS has a role in NMD regulation of mRNAs translated at the ER.

### UPF1 is present at the ER

The involvement of the ER-associated factor NBAS in the NMD response, together with its co-regulation of RNA targets with UPF1, led us to probe whether UPF1 localizes to the ER and interacts with NBAS. It has been previously reported that UPF1 localizes to the cytoplasm both in yeast (Atkin et al., 1995) and human cells (Applequist et al., 1997;Serin et al., 2001). There is, however, some evidence that UPF1 is also localized at the ER. First, UPF1 was found associated with cytoplasmic, but also with ER-bound polysomes (Jagannathan et al., 2014a). Moreover, a large-scale protein-protein interactome revealed that UPF1 interacts with components of the Syntaxin 18 complex; ZW10, p31 and STX18, where NBAS also resides (Aoki et al., 2009; Brannan et al., 2016).

Immunofluorescence analysis of HeLa cells transiently transfected with FLAG-UPF1 showed that UPF1 localizes to the cytoplasm, as previously suggested. However, upon incubation of HeLa cells with Digitonin, which permeabilizes the plasma membrane and consequently releases cytosolic components that are not anchored to cellular membranes, we observed a population of UPF1 that was resistant to Digitonin treatment and co-localized with the ER marker calnexin, indicating ER membrane association (Figure 4A). We next used the Proximity Ligation Assay (PLA) to probe for interactions of UPF1 and NBAS, with Sec61β, a component of the SEC61 channel-forming translocon complex that is a central component of the protein translocation apparatus at the ER membrane (Hartmann et al., 1994). PLA has been extensively used to detect interactions of many cellular proteins, including the core NMD factors UPF1 and UPF2 (Tatsuno et al., 2016). We detected a robust PLA interaction between UPF1 and Sec61β in HeLa cells, indicating that UPF1 localizes at the site of mRNA translation at the ER (Figure 4B). Next, we tagged endogenous NBAS with eGFP-3xFLAG-tag at its N-terminus, and performed PLA, which revealed that NBAS is also co-localized with Sec61β (Figure S3A). Altogether, these results show that both UPF1 and NBAS are present at the translocon, bringing them into close proximity with mRNAs being translated at the ER. Even though UPF1 is the core factor essential for NMD function, it is likely that other NMD factors will, at times, also partially localize to the ER. Previously, immunofluorescence was used to colocalize SMG6 and UPF3B with GRP78, a member of the heat shock protein 70 (HSP70) family that is present in the lumen of the ER (Sakaki et al., 2012). More recently, a large human interactome study identified UPF3B, in complex with UPF2 and UPF1, to be associated with SEC61A1, a component of the ER translocon (Hein et al., 2015). To investigate whether other components of the NMD pathway with previously established cytoplasmic localization are also present at the ER, we probed for the co-localization of UPF2 with Sec61β by PLA (Figure S3B). We observed a modest PLA signal indicating that a fraction of UPF2 is also associated with the translocon at the ER in HeLa cells. This modest co-localization of UPF2 with the ER is consistent with our previous observation that UPF2 plays only a minor role in the regulation of ER-associated genes.

**Figure 4.**
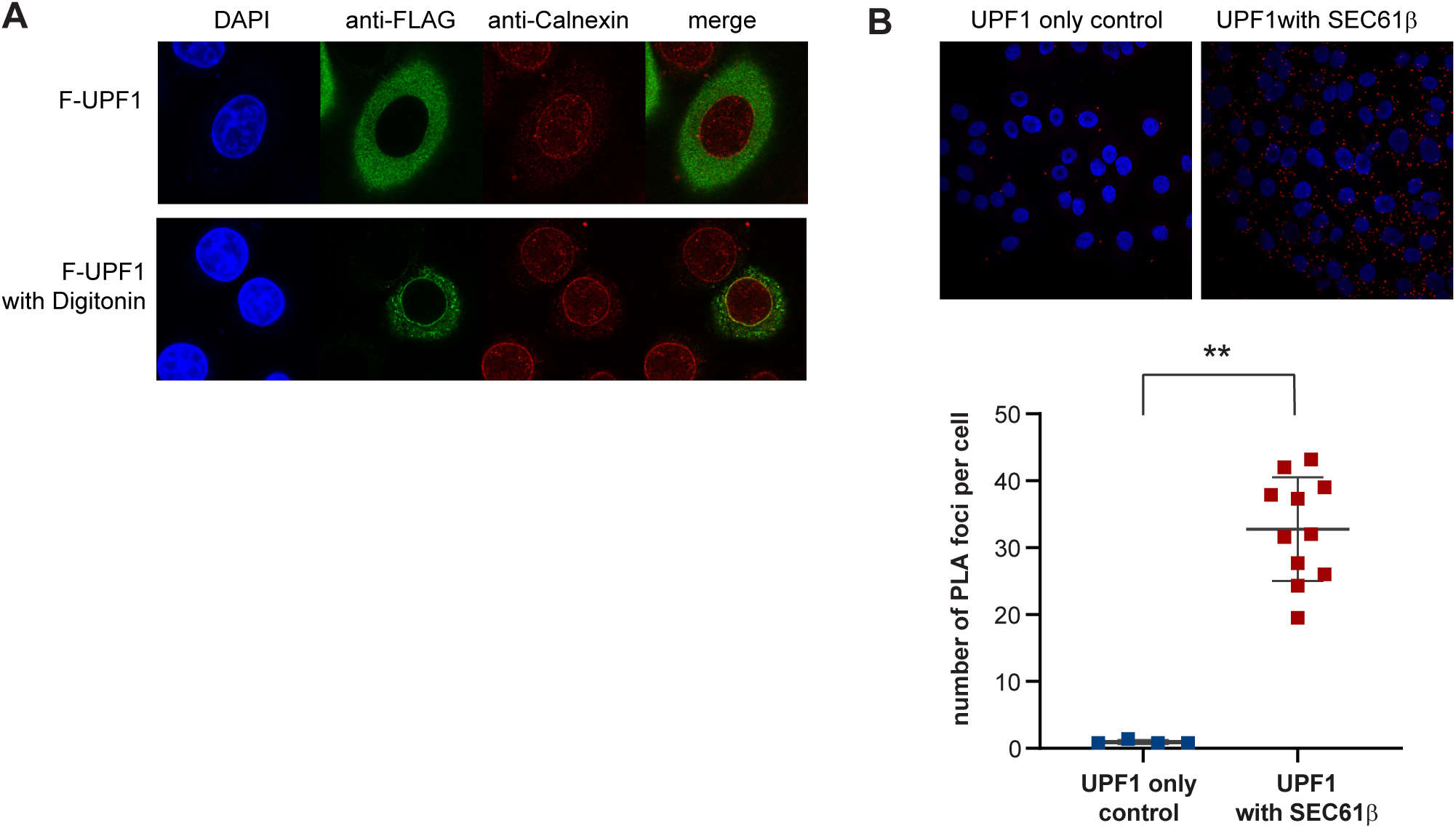
UPF1 localizes at the ER. (A) Immunofluorescence of HeLa cells transiently expressing FLAG-tagged UPF1 (in green) together with the ER-marker Calnexin (in red). Cell nuclei were visualized by DAPI staining. UPF1 predominantly shows a diffused cytoplasmic localization (top panel). Partial cell permeabilization with Digitonin revealed that a fraction of UPF1 is anchored at the ER membrane and co-localizes with Calnexin (bottom panel). (B) UPF1 is localized in the close proximity of the SEC61β translocon component at the ER. Proximity Ligation Assay (PLA) using antibodies against endogenous UPF1 and SEC61β proteins generated a discrete signal (red spots), indicating that the proteins are less than 40nm apart. The graph shows the quantification of PLA signal. Each point represents mean PLA count per cell in one captured frame. Significance was determined by two-tailed Mann-Whitney test: **: p<0.005. Data showing the co-localization of NBAS and UPF2 with SEC61β can be found in Figure S3.

### NBAS interacts with UPF1 and preferentially associates with the SURF complex

The presence of NBAS and UPF1 in proximity with the ER translocon, together with the co-regulation of RNA targets by NBAS and UPF1 as part of the ER-NMD response strongly suggested an interaction of these two NMD factors. To investigate this, we first tested for the co-purification of NBAS with UPF1 by co-immunoprecipitation in HeLa cells, in the presence or absence of RNases. We found that FLAG-tagged UPF1 co-immunoprecipitated with co-transfected T7-NBAS (Figure S4A). Since UPF1 phosphorylation is a later step in NMD activation it can be used as a diagnostic tool to infer the timing of recruitment of a particular protein to the NMD complex (Kashima et al., 2006). We used two UPF1 mutants that resemble the hypophosphorylated state of UPF1 when present in the early surveillance (SURF) complex (C126S), or the hyperphosphorylated UPF1 present in the late decay-inducing (DECID) complex (K498A) (Kashima et al., 2006; Weng et al., 1996)). As previously observed, the C126S mutation blocks UPF1 interaction with UPF2, yet it binds to T7-tagged NBAS (Figure S4B), whereas the K498A UPF1 mutant, displayed no binding (Figure S4B). This strongly suggests that NBAS is preferentially associated with the initial surveillance SURF complex, where UPF1 is hypophosphorylated. Interestingly, we previously observed similar results for the RNA helicase DHX34 (Hug and Cáceres, 2014), suggesting that most of the regulatory steps of the NMD pathway occur in the early stages of the NMD response. The observed physical interaction of NBAS with UPF1 was not RNA-dependent (Figure S4A). Nevertheless, we wanted to investigate the possibility that NBAS directly recruits NMD targets for degradation. We tested whether NBAS directly binds to mRNA using an mRNA capture assay, which relies on *in-situ* UV crosslinking, followed by affinity selection of mRNPs by oligo-dT cellulose (Piñol-Roma and Dreyfuss, 1992; Sanford et al., 2005). Affinity selection of mRNPs by oligo-dT cellulose showed that NBAS binds to mRNA (Figure S4C), opening the possibility that RNA-binding by NBAS contributes to selection of NMD targets at the ER.

Next, we used Förster Resonance Energy Transfer (FRET) to probe the interaction of NBAS and UPF1 in HeLa cells. This approach is based on the transfer of energy from the excited state of a donor fluorophore to an adjacent acceptor fluorophore when the two molecules are in the correct orientation and less than 10nm apart. FRET was detected by a reduction in the amount of energy that the donor releases as fluorescence, measured by Fluorescence Lifetime Imaging Microscopy (FLIM) ((Day et al., 2001; Ellis et al., 2008; Wouters et al., 2001). Co-transfection of GFP-NBAS and mCherry-UPF1 resulted in a reduction of the average donor fluorescence lifetime and an increase in the FRET efficiency, as compared to transfection of GFP-NBAS and mCherry (Figure 5A). This experiment strongly suggests that NBAS and UPF1 interact directly. To corroborate FRET results *in vivo*, in near-physiological concentrations of fluorescently labeled NBAS and UPF1, we performed Fluorescence Cross-Correlation Spectroscopy (FCCS) experiments. The cross-correlation signal is a direct indication of both molecules moving together: low amplitude of cross-correlation signal indicates that the labeled molecules diffuse separately, whereas high amplitude of cross-correlation signal is only achieved when both molecules are bound and diffuse together. The high cross-correlation signal observed in HeLa cells transiently expressing GFP-NBAS and mCherry-UPF1 indicated that NBAS and UPF1 interact directly (Figure 5B, black curve), in contrast to the low amplitude of cross-correlation signal observed in cells expressing GFP-NBAS with mCherry, as a negative control (Figure 5B, grey curve). Finally, we confirmed the interaction of endogenous NBAS and UPF1 using PLA. We observed a prominent PLA signal using antibodies against endogenous UPF1 and NBAS proteins, which was absent in cells that do not express NBAS (NBAS KO) (Figure 5C). These results robustly show that NBAS and UPF1 interact directly *in situ*.

**Figure 5.**
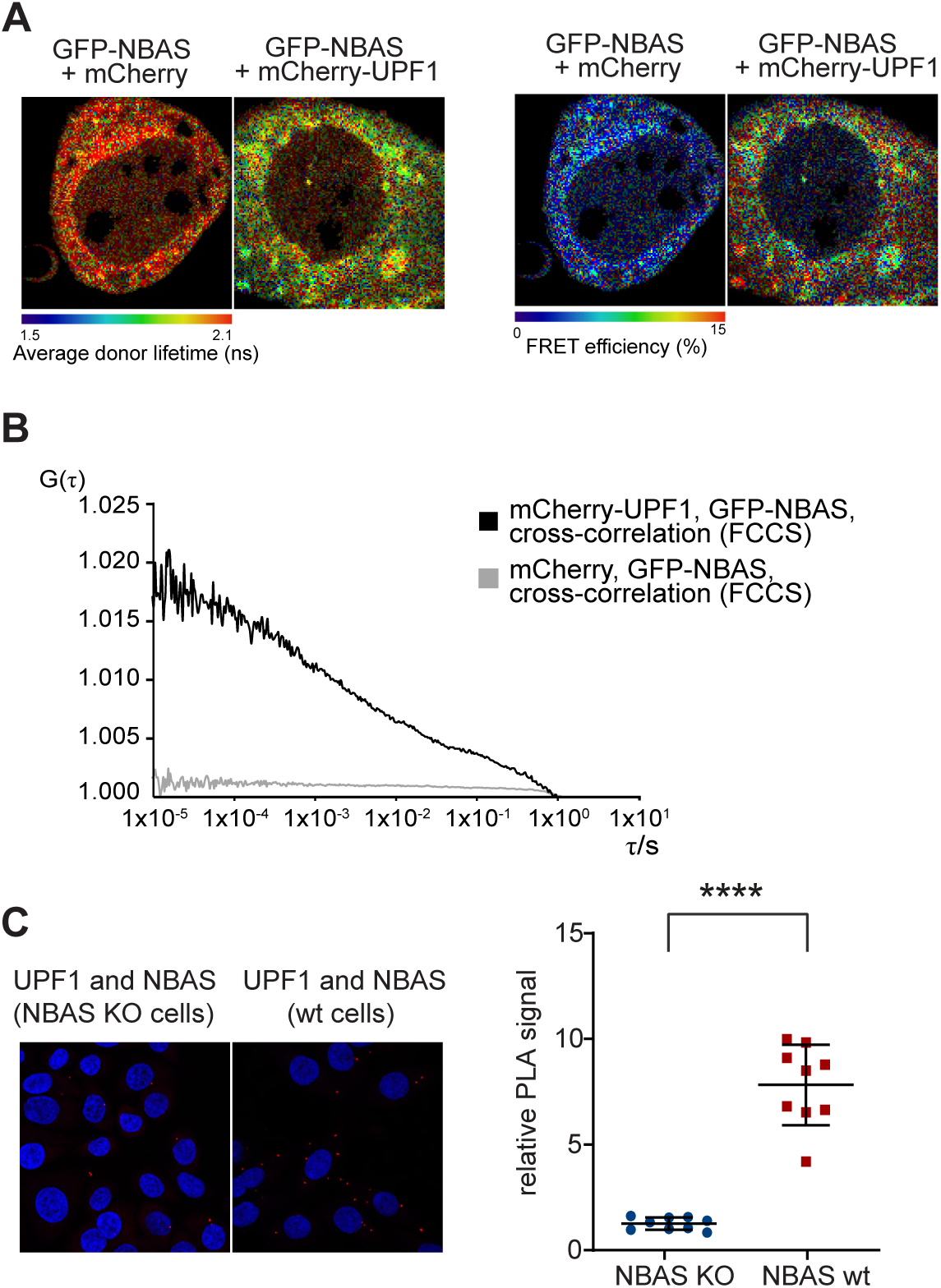
NBAS and UPF1 interact directly. (A) The interaction between NBAS and UPF1 was measured by FRET-FLIM. The average fluorescence lifetime of GFP-NBAS donor molecules was measured in HeLa cells expressing GFP-tagged NBAS together with mCherry-UPF1 or mCherry, as a control. Cells were pseudo-colored by average lifetime (ns) where red represents longer lifetime and green corresponds to a shorter lifetime. The co-expression of mCherry-UPF1 decreased the average fluorescence lifetime of GFP-NBAS in comparison to the mCherry control (left panel). Average fluorescence lifetime was used to calculate the FRET efficiency (right panel). Cells were pseudo-colored by FRET efficiency (%). (B) The interaction between NBAS and UPF1 was measured by Fluorescence Cross-Correlation Spectroscopy (FCCS) in HeLa cells transiently expressing GFP-NBAS and mCherry-UPF1 or GFP-NBAS and mCherry, as a control. Increased G(τ) amplitude of the average cross-correlation curves corresponds to higher cross-correlation (higher fraction of co-diffusing molecules) of GFP-NBAS and mCherry-UPF1 (in black), as compared to GFP-NBAS and mCherry average cross-correlation (in grey). (C) A direct interaction between endogenous NBAS and UPF1 was determined by PLA using antibodies against endogenous UPF1 and NBAS proteins in HeLa cells. The PLA signal in wild-type HeLa cells was compared to a HeLa NBAS knock-out (KO) cells, as a negative control. The PLA signal was quantified in the graph where each point represents mean PLA count in one captured frame, relative to NBAS KO negative control. Significance was determined by two-tailed Mann-Whitney test: ****: p<0.0001. Data showing the biochemical interaction of NBAS and UPF1 can be found in Figure S4.

### NBAS recruits UPF1 to the membrane of the ER

To probe the functional consequences of the NBAS-UPF1 interaction *in vivo* we used Fluorescence Correlation Spectroscopy (FCS). FCS is a non-invasive method with single-molecule sensitivity that allows the analysis of the dynamic behavior of fluorescent molecules with high temporal resolution and at low, physiologically-relevant concentrations in live cells (Bacia and Schwille, 2007; Liu et al., 2015; Papadopoulos et al., 2019). When the recorded fluorescence intensity fluctuations are caused by molecular movement, FCS measurements can be used to measure molecular mobility/diffusion rate of fluorescent molecules in a sub-femtoliter detection volume in live cells. Thus, free, fast diffusing molecular movement is reflected by autocorrelation curves that display decays in shorter characteristic times (Figure 6A, green curve), whereas bound or slow diffusing molecules are characterized by autocorrelation curves with slower decay times (Figure 6A, orange curve). We performed FCS measurements to derive the mobility of transiently transfected mCherry-UPF1, or mCherry control at the ER, in the perinuclear region of the cell, or in the cytoplasm at the cell periphery, in HeLa cells (Figure 6A, right panel). We observed that the mobility of mCherry-UPF1 in the cell periphery was very similar, irrespective of the levels of NBAS protein (Figure 6B; compare si-control, mCherry-UPF1, periphery (in gray) with si-NBAS, mCherry-UPF1, periphery (in black). By contrast, a significantly slower mobility of mCherry-UPF1was observed at the ER in control cells bearing physiological levels of NBAS, as reflected by a marked shift of the autocorrelation curve towards longer characteristic times (Figure 6B, red line). Depletion of NBAS increased the mobility of mCherry-UPF1 at the ER (Figure 6B, blue line), so that the diffusion time of mCherry-UPF1 was uniform within the cell. Fitting of the autocorrelation curves with a two-component diffusion model revealed that the characteristic decay time τ_D2_ of mCherry-UPF1 was considerably longer at the ER in the presence of NBAS (Figure 6C, in red), whereas NBAS depletion caused significant reduction of the τ_D2_ mCherry-UPF1 decay times (Figure 6C, in blue). The τ_D2_ decay time of mCherry-UPF1 was markedly shorter at the cell periphery than at the ER and was not affected by the depletion of NBAS (Figure 6C). The observed reduced mobility of mCherry-UPF1 at the ER strongly suggests that UPF1 is part of a slower diffusing protein complex, and that NBAS is required for the retention of UPF1 at the ER. In agreement, overexpression of NBAS led to even slower mCherry-UPF1 mobility at the ER (Figure 6D; compare wt NBAS level, mCherry-UPF1, ER (in black) with overexpressed GFP-NBAS, mCherry-UPF1, ER (in purple)). In accordance, the characteristic decay time τ_D2_ of mCherry-UPF1 was significantly increased by the addition of GFP-NBAS (Figure 6E). These results show that ER-localized NBAS acts to recruit UPF1 to the ER to activate a localized NMD response.

**Figure 6.**
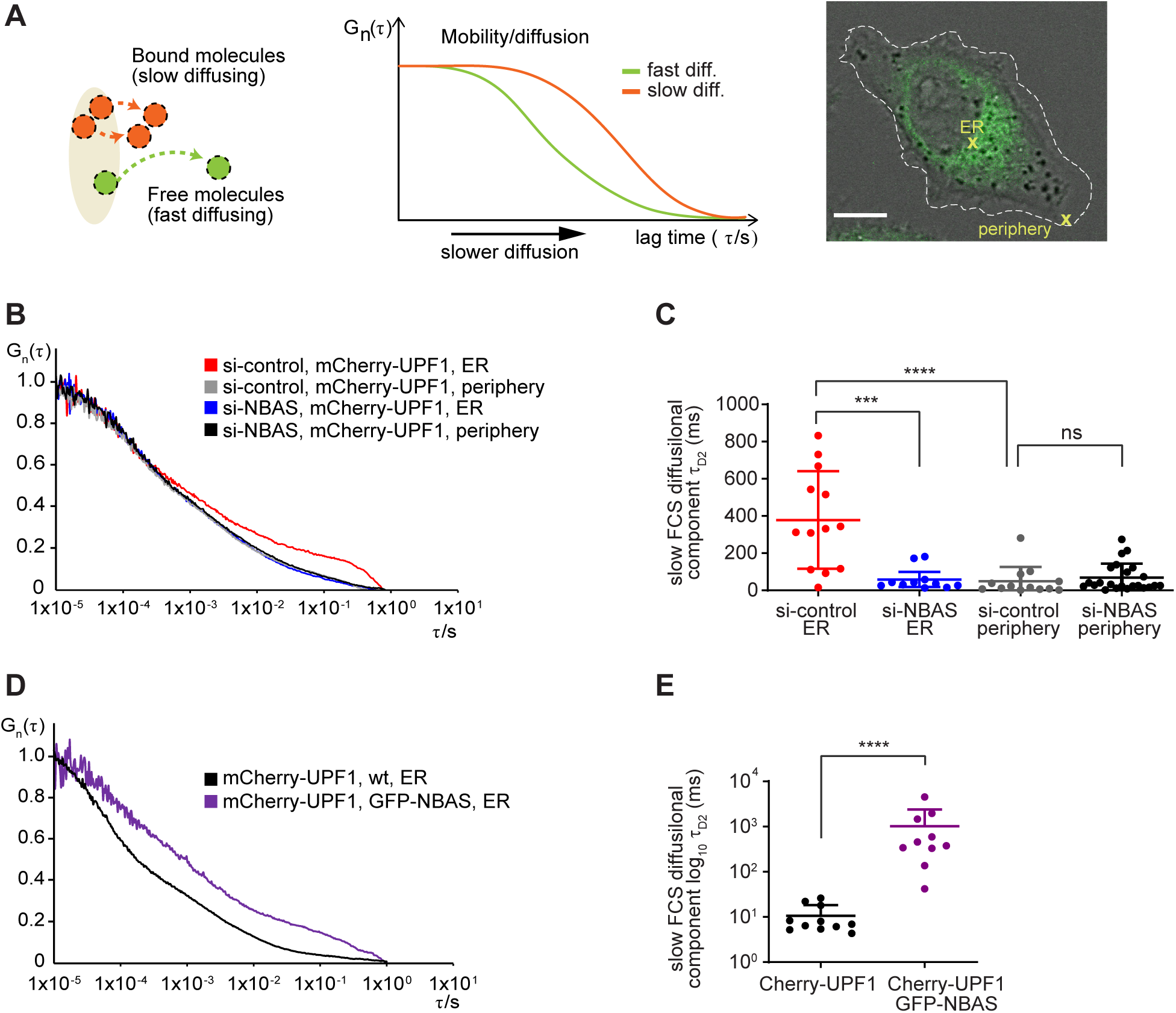
NBAS recruits UPF1 to the ER. (A) Schematic representation of the principle of Fluorescence Correlation Spectroscopy (FCS). See text in methods section for details. (B) Average autocorrelation curves of mCherry-UPF1 at the ER or cell periphery in the presence (si-control) or absence of NBAS (si-NBAS) are displayed. (C) Graph showing the derived characteristic decay times τ_D2_ (slow FCS component) of mCherry-UPF1, after fitting of the autocorrelation curves with a two-component model for diffusion (plus triplet correction). Significance was determined by the two-tailed Mann-Whitney test: ****: p<0.0001; ***: p<0.001; ns: not significant. (D) Average autocorrelation curves of mCherry-UPF1 at the ER with physiological (wt) or increased (GFP-NBAS) level of NBAS expression. (E) Characteristic decay times τ_D2_ (slow FCS component), derived as in (C) of mCherry-UPF1 at the ER. Significance was determined by the two-tailed Mann-Whitney test: ****: p<0.0001. Data showing FCS control experiments can be found in Figure S5.

## DISCUSSION

### An NMD response at the ER

The targeting of mRNAs to specific subcellular sites for local translation has an important role in many cellular processes and during development (Decker and Parker, 2006). In particular, translation of secreted and integral membrane proteins occurs on ER-bound ribosomes, whereas cytosolic protein synthesis mostly occurs on ribosomes dispersed throughout the cytoplasm. A subset of transcripts that are targeted for translation at the ER encode a leading signal peptide that is recognized after emerging from the ribosome by the signal recognition particle (SRP). These SRP-bound transcripts are subsequently targeted to the ER, and the resulting translated peptides are translocated across or inserted into the membrane by the Sec61 translocation channel (Voorhees and Hegde, 2016; Walter and Johnson, 1994). However, recent evidence showed that mRNAs encoding cytosolic proteins can also be translated by ER-associated ribosomes, suggesting a more diverse role for the ER in mRNA translation (Jagannathan et al., 2014b; Reid and Nicchitta, 2015). Thus, despite the ER representing a specialized environment for the translation of a large proportion of cellular mRNAs, it remains unclear how RNA quality control pathways in general, and NMD in particular, operate on those mRNAs. We speculate that transcripts coding for secreted and integral membrane proteins will not have sufficient exposure to the NMD quality control in the cytoplasm due to the intrinsic nature of ER-associated translation. This is spatially and temporally distinct from mRNA translation in the cytosol, as shown by the fact that ER-targeted transcripts are only translationally active once they encounter the ER membrane (Wu et al., 2016). Since NMD is dependent on active translation, we hypothesize that ER-targeted transcripts will fail to undergo NMD in the cytoplasm, until their translation is activated at the ER membrane. We propose that these transcripts are instead targeted for degradation by an ER-NMD dedicated pathway that is responsible for the quality control of cellular mRNAs that are translated at the ER.

Here, we have identified such a localized NMD response at the ER, which relies on the activity of NBAS to recruit UPF1 and co-regulate the stability of target RNAs, which are preferentially enriched for mRNAs localized and translated at the ER (Figures 2 and S2). Using smRNA FISH we were able to visualize the site of this NBAS-dependent NMD response of mRNAs translated at the ER (Figure 3). We observed that UPF1 interacts with the translocon component Sec61β at the ER membrane, as was also found for NBAS (Figures 4 and S3). Use of FRET-FLIM, FCCS and PLA confirmed that NBAS and UPF1 interact directly (Figure 5). In agreement with this, an interactome study, using a UPF1-HaloTag fusion protein pull-down followed by quantitative mass spectrometry, revealed interactions of the core NMD factor UPF1 with components of the Syntaxin-18 complex, including Sly1, STx10, p31 and ZW10 (Brannan et al., 2016). Together with our observation that SEC13 (a component of COPII vesicles involved in ER transport) is also a conserved NMD factor (Casadio et al., 2015), this supports the model of a functional link between NMD and ER vesicular transport and/or ER homeostasis. These results, together with evidence provided by FCS experiments demonstrates that NBAS, which is anchored at the membrane of the ER, promotes the recruitment of the core NMD factor UPF1 and thereby activates the NMD response at the ER (Figure 6). We postulate the existence of a dedicated branch of the NMD pathway that acts on transcripts localized and translated at the ER, which we term ER-NMD.

### Biological role of ER-NMD pathway

Recent evidence supports a role for NMD in modulating the ER stress response by ensuring appropriate activation of the Unfolded Protein Response (UPR) (Goetz and Wilkinson, 2017). This pathway is able to sense and respond to excessive amounts of misfolded proteins in the ER (Walter and Ron, 2011). Interestingly, NMD targets mRNAs encoding several UPR components, including the UPR sensor IRE1α, as well as ATF-4 and CHOP that are activated by PERK branch signalling. Thus, NMD can act to control the threshold of cellular stress that is necessary to activate the UPR (Gardner, 2008; Karam et al., 2015; Sieber et al., 2016). As a consequence, appropriate level of NMD activity is required to protect cells from detrimental stress response activation. In particular, pathological UPR activation is recognized as a major causal mechanism in neurodegenerative diseases including Alzheimer’s, Parkinson’s and prion diseases, which are associated with the accumulation of misfolded disease-specific proteins (Hetz and Mollereau, 2014). A chronic activation of PERK/eIF2α-P signaling results in a sustained reduction of global protein synthesis rates in neurons, caused by the phosphorylation and inactivation of eIF2α, which blocks translation at the level of initiation. Inappropriate eIF2α phosphorylation contributes to disease pathogenesis, and is observed in patient’s brains, and in mouse models of protein-misfolding neurodegenerative diseases (Freeman and Mallucci, 2016; Hoozemans et al., 2009; Moreno et al., 2012). Thus, limiting the chronic activation of the UPR could have a neuroprotective effect in neurodegenerative diseases.

We hypothesize that modulation of the activity of the ER-NMD pathway could be crucial to regulate the response to ER stress. It is tempting to speculate that increasing the activity of the ER-NMD pathway will have a neuroprotective effect by limiting excessive UPR signalling in neurodegeneration. Indeed, a protective role for UPF1 has already been shown in primary neuronal models of amyotrophic lateral sclerosis (ALS) and frontotemporal dementia (FTD) induced by overexpression of the RNA-binding proteins, TDP43 and FUS (Barmada et al., 2015). Moreover, NMD has been shown to protect against the effects of hexanucleotide repeat expansion in C9orf72 (C9-HRE) found in *Drosophila* and cellular models of ALS/FTD (Ortega et al., 2020; Xu et al., 2019)

In summary, we have uncovered a localized NMD response at the ER, which provides spatial and temporal separation from cytoplasmic NMD acting on mRNAs that are translated at the ER. This could have important implications to protect cells from ER stress/dysfunction that results from the accumulation of aberrantly processed RNAs, leading to the production of truncated proteins and affecting cellular homeostasis. This could be of particular relevance for many disease processes, such as neurodegenerative diseases, where an exacerbated ER stress is observed. Future studies will aim to further characterize this novel ER-NMD pathway; investigating its physiological role, as well as the biological consequences of manipulating its activity.

## SUPPLEMENTAL INFORMATION

Supplemental information includes 5 figures and 4 tables.

## ACKNOWLEDGEMENTS

We are grateful to Andrew Peden (University of Sheffield) for the gift of HeLa C1 cells, a GFP-based secretion reporter and for advice on secretion experiments, and Konstantin Lukyanov (Moscow) for the gift of an NMD+ reporter. We thank Matthew Pearson (Advanced Imaging), Elisabeth Freyer (Flow Cytometry), and Craig Nicol (Graphics) from the MRC IGMM core support facilities for technical assistance and advice, Shelagh Boyle for her help with the smRNA FISH set up, and Alison Dun and Rory Duncan at the MRC-funded Edinburgh Super-Resolution Imaging Consortium (ESRIC) for both expertise and infrastructure. We are thankful to our MRC HGU colleagues, Andrew Jackson, Ian Adams and Wendy Bickmore for helpful discussions and critical reading of the manuscript. This work was supported by a Chancellor’s fellowship of the University of Edinburgh (D.K.P) and an Institutional Strategic Research Fund (ISSF3) (J-J.S.) and core funding to the MRC Human Genetics Unit from the Medical Research Council (J.F.C).

## AUTHOR CONTRIBUTIONS

D.L. and J.F.C. conceived and designed the project. D.L. performed secretion assays, NMD assays, RNA FISH, cell biological, FRET-FLIM and FCS experiments. K.J.J. and M.M.M. carried out RNA-seq profiling of NBAS and UPF1 knock-down cells, K.J.J. generated the endogenously tagged NBAS cell line and together with R.S.Y and M.S.T performed all bioinformatic analysis. L.M. performed the analysis of FISH signal distribution. D.K.P. and J.J.S. performed, analyzed and interpreted the FCS and FCCS experiments. The manuscript was co-written by all authors.

## LEAD CONTACT AND MATERIALS AVAILABILITY

Further information and requests for resources and reagents should be directed to and will be fulfilled by the Lead Contact, Javier F. Caceres

## Materials and Methods

### Cell Culture

HeLa and DLD-1 cell lines were maintained at 37°C in the presence of 5% CO_2_, in DMEM media with high glucose, GlutaMAX™ Supplement, pyruvate (Gibco Life technologies; 10569010) supplemented with 10% FCS. HeLa C1 cells and 3xFLAG-eGFP-NBAS cells were maintained in the same media with the addition of 1µg/ml of Puromycin.

### Cell Transfections

Cells were grown without antibiotic prior to transfections. All transfections were carried out in Opti-MEM reduced serum medium (Gibco, 31985047). Transfections of siRNA oligos was done using DharmaFECT 1 (Dharmacon, T-2001-03) following manufacturer’s protocol. Plasmids were transfected using Lipofectamine 2000 (ThermoFisher Scientific, 11668019) following manufacturer’s instructions. Lipofectamine 2000 was also used for the co-transfection of plasmids and siRNA oligos. For depletions, cells were plated in 12-well plates and transfected with 30pmol of indicated siRNAs. Cells were then expanded into 6-well plates and transfected with 50pmol of the same siRNAs on the third day. Cells were harvested for analysis on day 5 after the first depletion. For the fluorescence based NMD assays HeLa cells were treated as above, with the addition of 50ng NMD+ reporter that was co-transfected with siRNAs during the second round of depletion. For the NMD assays cells were no more than 80% confluent at any stage of the experiment. For total RNA-sequencing and subcellular fractionation, cells were plated in 6-well plates and transfected with 30pmol of indicated siRNAs. Following day, cells were expanded into 10cm plates and transfected with 30pmol of the same siRNAs on the third day. Cells were harvested for analysis on day 4 after the first depletion. For co-immunoprecipitation, 80% confluent HeLa cells in 10cm dishes were transfected with 5μg of each T7-tagged and FLAG-tagged expression vectors for 24-48 h. For Immunofluorescence and FRET-FLIM interaction measurements 200ng of GFP-NBAS was co-transfected with 100ng of mCherry plasmids into HeLa cells in 6-well plates. 24 h after transfection cells were plated onto coverslips and fixed next day.

### Flow cytometry-based assay for measuring constitutive secretion

The secretion assay was performed as previously described (Gordon et al., 2010). HeLa C1 cell mock-depleted or depleted of UPF1, UPF2, DHX34, SEC13, NBAS, p31, RINT, ZW10 and STX18 were incubated with 1µM D/D Solubilizer (ligand) for 1.5h to facilitate secretion or left untreated for comparison. Cells were trypsinized, washed with cold PBS and placed on ice to stop further secretion. To measure the amount of GFP reporter remaining, cells were analysed by FACS BD AccuriTM (488nm excitation laser, FL-1: 533/30nm emission filter) running the software BD AccuriTM C6. Gates were set using a non-fluorescent control. GFP expression was analysed by gating the single cell population in SSC-H/SSC-A dotplot, followed by debris-exclusion gate in SSC-A/FSC-A dotplot. Between 5000 and 10000 single cells were analysed for each sample. The mean GFP fluorescence was calculated with FlowJo™ Software (Version 10.6.0.). The relative GFP fluorescence remaining after secretion was calculated as a ratio between the mean fluorescence of depleted cells incubated with the ligand and depleted cells. The relative fluorescence remaining of untreated C1 cells was set to 1, and the threshold for a secretion defect was set to two.

### NMD reporter assays

HeLa cells stably expressing HBB wt or HBB NS39 reporter were mock-depleted or depleted twice of UPF2, NBAS, STX18 and STX5, and harvested 5 days after the first depletion, as described above. The expression of HBB reporters was analysed by qRT-PCR. Alternatively, HeLa HBB NS39/wt cells were treated with 2µg/ml Actinomycin D to block transcription and/or 2µg/ml Brefeldin A to block secretion for 6 hrs as indicated. The stability of HBB reporters was determined by qRT-PCR. The florescence-based NMD assay was performed as described previously (Pereverzev et al., 2015). HeLa cells mock-depleted or depleted of UPF2, NBAS, STX18 and STX5 expressing NMD+ reporter were harvested 5 days post depletion as described above. NMD activity was determined by the quantitative measurement of the red and green fluorescence using FACS BD LSR-Fortessa™ X-20 SORP with BD FACSDiva™ software. Gates were set using a non-fluorescent control. Compensation was applied in BD FACSDiva™ software using single transfected cells. Green fluorescence expression (488-525/50) was analysed by gating the single cell population in SSC-H/SSC-A scatterplot, followed by debris-exclusion gate in SSC-A/FSC-A scatterplot, followed by gating only the red fluorescent positive cells in 405-450/50 (autofluorescence) and 561-610/20 dot plot. Gate settings were kept constant during the experiment. The data was analysed with FlowJo™ Software (Version 10.6.0.). Between 4500 and 12000 red fluorescent cells were analyzed in each biological replica. NMD disruption was determined by the increase of green fluorescence in red cells, normalized to mock-depleted cells.

### Quantitative RT-PCR

Total RNA was isolated using RNeasy Mini kit and resuspended in nuclease-free water at 100ng/µl. qRT–PCR was performed using SuperScriptIII One-Step RT–PCR Kit (Invitrogen) following the manufacturer’s instructions. All RT-PCRs were run on the CFX96 Real-Time System (Bio-RAD machine, following this program: RT at 50°C for 30min, 95°C for 2min, then 40 cycles of 95°C for 30sec, 55°C for 20sec, 70°C for 20sec followed by the plate read step. Each sample was run in 3 technical replicates. Primers were designed using Roche Real-Time Ready Configurator and combined with Roche Universal Probe Library (See Table S4). Gene expression data was analysed by the delta Ct method, with each gene normalised to housekeeping gene *POL2RJ*. Unpaired two-tailed t-test was used for statistical analysis.

### Gene expression profiling: RNA extraction, library preparation and RNA-sequencing

HeLa cells were depleted as described above. Total RNA was isolated from depleted cells by phenol-chloroform extraction and treated with TURBO DNA-free™ DNase I kit (Invitrogen Ambion; AM1907). Libraries were prepared following NEBNext Ultra Directional RNA Library Prep Kit for Illumina (New England Biolabs; #E7420) and Agencourt AMPure XP Beads (Beckman Coulter; A63881). Samples were ligated to barcode primers 1-20 of NEBNext Index Primers for Illumina sets 1 and 2 (New England Biolabs) and libraries analysed using DNA High Sensitivity chip on an Agilent 2100 Bioanalyzer before being pooled. 150 base pair, paired-end sequencing was performed using the S2 flow cell on the NovaSeq 6000 System (Illumina Inc.). Library molarity for sequencing was calculated using Qubit dsDNA quantification results and fragment size information from Bioanalyzer results. Sequencing was performed by Edinburgh Genomics, Edinburgh, UK.

### Cell fractionation

HeLa cells were mock-depleted or depleted twice of UPF1, UPF2 and NBAS as described above. Cellular fractionation was performed as described before (Jagannathan et al., 2011) with some modifications. Briefly, four days after the first depletion, cells were detached using trypsin and washed twice in 1ml of ice-cold PBS (500g, 10 minutes, 4°C). Cellular pellets were resuspended in 0.4 ml of permeabilization buffer (110mM KOAc, 25mM K-HEPES pH 7. 2, 2.5mM Mg(OAc)_2_, 1mM EGTA, 0.015% digitonin, 1mM DTT, 1× Complete Protease Inhibitor Cocktail, 40U/mL RNaseOUT™) and incubated for 5 minutes on a rotating wheel at 4°C. The resulting cytosolic fractions were recovered by centrifugation at 2000g for 10 minutes, 4°C. The cells were then lysed in 0.4 ml of NP-40 lysis buffer (400mM KOAc, 25mM K-HEPES pH 7.2, 15mM Mg(OAc)_2_, 1% (v/v) NP-40, 1mM DTT, 1× Complete Protease Inhibitor Cocktail, 40U/mL RNaseOut) for 30 minutes on ice. Membrane fraction was recovered by centrifugation at 7000g for 10 minutes at 4°C. Both cytosolic and membrane fractions were clarified by centrifugation at 7500g for 10 minutes at 4°C. Digitonin, DTT, Complete Protease Inhibitor Cocktail and RNaseOUT™ were added fresh to the buffers. RNA from both fractions was isolated using PureLink RNA Mini Kit (Life Technologies) according to manufacturer’s instructions. DNA was removed using TURBO DNA-free™ DNase I kit (Invitrogen Ambion; AM1907)

### RNA-sequencing analysis

FASTQ files were quality control checked for base and sequence quality scores, and adapter contamination using fastQC (v0.11.7; Babraham Bioninformatics). Reads were aligned using Spliced Transcripts Alignment to a Reference (STAR, v2.5.1b) (Dobin et al., 2013)) or pseudoaligned using kallisto (v0.43.1; (Bray et al., 2016)). Kallisto index was created by combining “all basic gene annotation” and “long non-coding RNA gene annotation” fasta files, downloaded from Gencode (www.gencodegenes.org/releases/current.html), Release 27 (GRCh38.p10). Kallisto was run with 100 bootstraps. Reads per feature in STAR output files was counted using HTSeq (v0.9.1; Anders et al., 2015). Abundance files were then analysed for fold changes by running DESeq2 (v1.14.1) differential expression analysis. Gene Ontology enRIchment anaLysis and visuaLizAtion (GOrilla) (http://cbl-gorilla.cs.technion.ac.il/) (Eden et al., 2009) and Database for Annotation, Visualization and Integrated Discovery (DAVID) (https://david.ncifcrf.gov/home.jsp) (Huang et al., 2009) were used for Gene Ontology (GO) analysis. To investigate genes of specific GO terms, genes were annotation using biomaRt Bioconductor R package (Durinck et al., 2005). Overlap with experimental ER-localisation was carried out by downloading relevant supplementary data. APEX-seq (Fazal et al., 2019), ER Fractionation (Reid and Nicchitta, 2012), ER Ribosome Profiling (Jan et al., 2014). Gene names converted using DAVID (https://david.ncifcrf.gov/home.jsp)(Huang et al., 2009), 1.5-fold enrichment at ER cut off applied. For fractionated RNA-sequencing, gene expression in both fractions in control cells was compared by using variant stabilised transformation of tpm, averaged across replicates. Membrane-associated genes were defined as those >2-fold higher expression in membrane fraction than cytoplasmic fraction, all other genes were termed non-membrane. Membrane-enrichment was confirmed by overlap with three experimental ER datasets (see above). Changes in gene expression resulting from depletion of individual NMD factors was measured by percentage of ‘membrane-associated’ genes in membrane fraction, and of ‘non-membrane’ genes in cytoplasmic fraction regulated. Genes were determined as regulated if they were significantly (p<0.05) increased in expression when the relevant factor was depleted.

### Single-molecule (sm) RNA FISH

smRNA FISH was performed following the Stellaris RNA FISH protocol for adherent cells available online at www.biosearchtech.com/stellarisprotocols, using RNase-free reagents. Transcript-specific Quasar® 570-labeled Stellaris probes were designed with the Stellaris® RNA FISH Probe Designer (Biosearch Technologies, Inc., Petaluma, CA) available online at www.biosearchtech.com/stellarisdesigner, and resuspended in TE buffer to make 12.5µM Probes stock. The probe sequences are listed in the Table S4. HeLa cells were depleted as described above, with the addition of mEmerald-ER-3 plasmid (1µg per well in 6-well plates) during the second round of depletions. mEmerald-ER-3 was a gift from Michael Davidson (Addgene plasmid # 54082; http://n2t.net/addgene:54082; RRID:Addgene_54082). Cells were seeded on high precision coverslips (Marienfeld, #0107052), washed twice with PBS and fixed with 4% formaldehyde at room temperature for 10min. Cells were permeabilized in 70% ethanol for at least 1hr. Hybridization of probes was performed in a humidified chamber at 37°C for 15hrs. Probes hybridization and all subsequent steps were carried out in dark. After the hybridization, coverslips were washed as described in the protocol. Nuclei were counterstained with 5ng/ml of DAPI in buffer A for 30min. Cell outlines was determined by staining with the addition of 20nM MemBright 640nm dye (Collot et al., 2019) in buffer B for 15min at room temperature. Coverslips were mounted in Vectashield and sealed with nail polish.

### Image Capture and Analysis

For smRNA FISH experiment, epifluorescent images were acquired using a Photometrics Coolsnap HQ2 CCD camera and a Zeiss AxioImager A1 fluorescence microscope with a Plan Apochromat 100x 1.4NA objective, a Nikon Intensilight Mercury based light source (Nikon UK Ltd, Kingston-on-Thames, UK) and either Chroma #89014ET (3 colour) or #89000ET (4 colour) single excitation and emission filters (Chroma Technology Corp., Rockingham, VT) with the excitation and emission filters installed in Prior motorised filter wheels. A piezoelectrically driven objective mount (PIFOC model P-721, Physik Instrumente GmbH & Co, Karlsruhe) was used to control movement in the z dimension. Step size for z stacks was set at 0.2 µm. Hardware control, image capture and analysis were performed using Nikon Nis-Elements software (Nikon UK Ltd, Kingston-on-Thames, UK). Images were deconvolved using a calculated point spread function with the constrained iterative algorithm of Volocity (PerkinElmer Inc, Waltham MA). Image analysis was carried out using the FIJI/ImageJ software (2.0.0-rc-69/1.52p) (Schindelin et al., 2012). To measure the distribution of FISH signal within cells, a maximum intensity projection of each deconvolved z-stack was created and the DAPI signal was thresholded to create a mask of the nucleus. This mask was inverted and a euclidean distance map was created using the “Distance Map” function. This resulted in an image where all pixels that were inside the nucleus had a value of zero and the other pixels had values based on their distance from the closest nucleus edge (in pixels). To detect the centroids of FISH spots in the FISH signal channel, the ImageJ function “Find Maxima” was used with a prominence setting of 800. The x-y co-ordinates of each maximum point was measured on the distance map, giving a distance of the FISH spots to the edge of the nucleus. These values were converted to microns using the pixel spacing value of the original image. For immunofluorescence and PLA experiments images were acquired on a Nikon Confocal A1R confocal microscope using a Plan Apochromat 100x 1.4NA objective. The microscope comprises a Nikon Eclipse TiE inverted microscope with Perfect Focus System and is equipped with 405nm diode, 457/488/514nm Multiline Argon, 561nm DPSS and 638nm diode lasers. Detection is via four Photomultiplier tubes (2x standard Photomultiplier tubes and 2x GaAsP PMTs). Data were acquired using NIS Elements AR software (Nikon Instruments Europe, Netherlands). Z-stacks of images were acquired with a 0.2 µm step, scan size 1024×1024, 1.2x zoom and 2x frame averaging. Image analysis was carried out using the FIJI/ImageJ software.

### Design and screening of CRISPR cell lines

The design of the guide RNAs (gRNAs) was undertaken, as previously described (Ran et al., 2013). gRNAs were designed using sgRNA Designer CRISPRko (broad institute, https://portals.broadinstitute.org/gpp/public/analysis-tools/sgrna-design) and Cas-Designer (RGEN Tools, http://www.rgenome.net/cas-designer/). For the tagged-NBAS cell line, four gRNAs were selected by closest proximity to the start codon and highest predicted efficiency. For each guide RNA, the complementary sequence was determined and a BbsI restriction site added to both oligos. Designed gRNAs were ordered as custom single stranded DNA oligos with an extra G at the 5’ end, and 5’ phosphate at the reverse complement DNA strand (IDT). Top and bottom strands of gRNAs were annealed at a concentration of 100μM and cloned into the px459 V2.0 vector using BbsI restriction cloning. 1μl of a 1:250 dilution of annealed gRNAs was ligated with the T4 DNA Ligase (NEB) into 36ng of the px459 V2.0 vector. To assess cutting efficiency of gRNAs, each plasmid was transfected into HeLa cells and RNA extracted as above. PCR over the target region was performed with custom primers (Table S4) and the resulting products assessed by DNA electrophoresis. The PCR products were also sent for Sanger sequencing and the traces analysed by TIDE (REF) and ICE (REF) to give exact cutting efficiencies for each guide. Repair template was ordered as a custom plasmid from IDT on a pUCIDT-AMP backbone. For the generation of NBAS knock out (KO) HeLa cells, two guides (NBAS KO_A and NBAS KO_B) targeting the 5^th^ exon of the NBAS gene were cloned into the pSpCas9n(BB)-2A-GFP vector encoding the (D10A nickase mutant (PX461) as described above. pSpCas9n(BB)-2A-GFP (PX461) (Addgene plasmid # 48140; http://n2t.net/addgene:48140; RRID:Addgene_48140). The gRNA/Cas9 plasmid and repair template plasmid were transfected into HeLa cells as described above. After 24hr media was supplemented with 1.5ug/ml puromycin to select for cells expressing the gRNA/Cas9 plasmid. Once control cells were dead, surviving cells’ fluorescence was measured by FACS and single cells deposited into wells of a 96-well plate. After approx. 2 weeks clonal expansion, genomic DNA was extracted by lysing with DirectPCR Lysis Reagent Cell (Peqlab, VWR) supplemented with 0.5 μg/μl Proteinase K (Invitrogen), at 55°C O/N. PCR of the N-terminal locus of NBAS was performed using previously mentioned custom primers and SYBR green Master Mix (REF). PCR products were resolved on an agarose gel for genotyping. The correct repair template integration was validated by genomic DNA sequencing and expression of tagged-NBAS was validated by Western blotting. Screening for the NBAS KO clones was performed as above. HeLa cells co-transfected with NBAS KO_A and _B pSpCas9n(BB)-2A-GFP plasmids were FACS-sorted into 96-well plates 48h after transfection. Growing colonies were screened by PCR of genomic DNA as described, using NBAS KO F and R primers. NBAS expression was assessed by Western Blotting and genomic DNA of the NBAS KO was checked by sequencing.

### Immunoprecipitation and Western Blotting

Cells were washed and harvested in ice-cold PBS before pellets were lysed with immunoprecipitation (IP) buffer [20mM Tris-HCl pH 8, 150mM NaCl, 1mM EDTA, 1% NP-40, 0.2% NaDeoxycholate, Complete Protease Inhibitor (Roche), Phospho STOP (Roche), 1mMDTT] for 20min on ice. Cell lysates were treated with 40–80 mg/ml RNase A per 1 ml of extract. Lysates were precleared with Dynabeads Protein-G (Novex, Life Technologies) for 1hr, rotating at 4°C. Anti-FLAG M2 antibody coupled magnetic beads (Sigma) were washed x3 with IP buffer before incubation with the precleared lysate overnight rotating at 4oC. Beads were washed x5, each for 5min with IP buffer and then bound protein was eluted by boiling in SDS sample buffer supplemented with reducing agent for 5min. Proteins were resolved by SDS-PAGE using NuPAGE 3-8% Tris-Acetate precast gels (Novex, Life Technologies, Invitrogen) run for ∼1hr at 170V in 1x Tris-Acetate running buffer. Protein transfer was achieved using the iBlot™ 2 Gel Horizontal Transfer Device (Invitrogen). Nitrocellulose membranes were blocked in 5% BSA in PBS/Tween 20 (0.1%) for a minimum of 1hr at room temperature and probed with the appropriate primary antibody diluted in blocking solution 1:3000. FLAG antibody was used at 1:10,000 dilution and T7 antibody was used at 1:5,000 dilution. HRP-conjugated secondary antibodies (BioRAD) were used at 1:10,000 and blots developed with ChemiGlow detection reagent and visualized using ImageQuant LAS 4000 chemiluminescent camera (GE Healthcare).

### In situ UV cross-linking mRNP capture assay

In situ UV cross-linking mRNP capture protocol was adapted from (Piñol-Roma and Dreyfuss, 1992). Briefly, cells, grown in 150mm plates, were irradiated with 0.15Jcm^−2^ at 254-nm UV light, on ice, in ice-cold PBS. Cells were then scraped from plates in PBS and pelleted. Cells were lysed in 1ml of ice-cold lysis buffer (20mM Tris-HCl (pH 7.5), 500mM LiCl, 0.5% LiDS (wt/vol, stock 10%), 1mM EDTA and 5mM DTT), and then sonicated 30s on / 30s off for 5 cycles at 4°C. After 10mins incubation at 4°C, the lysates were fractionated by centrifuge (15,000 xg, 4°C, 15min), 10% sample was stored as input. 100µl per sample oligo(dT)25 magnetic beads were equilibrated in lysis buffer and incubated with lysate overnight at 4°C with rotation. Beads were washed once with 1ml of lysis buffer, twice with Buffer 1 (20 mM Tris-HCl (pH 7.5), 500 mM LiCl, 0.1% LiDS (wt/vol), 1 mM EDTA and 5 mM DTT), twice with Buffer 2 (20 mM Tris-HCl (pH 7.5), 500 mM LiCl, 1 mM EDTA and 5 mM DTT) and twice with Buffer 3 (20 mM Tris-HCl (pH 7.5), 200 mM LiCl, 1 mM EDTA and 5 mM DTT). All washes for 5mins at 4°C. Captured mRNPs were eluted from the beads with 100µl elution buffer (20 mM Tris-HCl (pH 7.5) and 1 mM EDTA) for 3min at 55°C and treated with 200U of RNase A in 500μl of 10× RNase buffer, for 1h at 37 °C. Liberated mRNA binding proteins were mixed 1:1 with SDS PAGE sample buffer (1x LDS sample buffer, 1x reducing agent), and resolved by 3-12% Tris-acetate SDS-PAGE and analyzed by Western blotting.

### Immunofluorescence

Cells were grown on coverslips, fixed with 4% paraformaldehyde at room temperature for 10min, washed with PBS and permeabilised with 0.5% Triton X-100 at room temperature for 10min. Coverslips were then incubated for 1 hour with block buffer (1% BSA, 0.01% Triton X-100 in PBS). Primary antibodies (diluted 1:500 in block buffer) were incubated with coverslips in humidified chamber overnight at 4°C. Coverslips were washed 3 times with wash buffer (0.01% Triton X-100 in PBS). Secondary antibodies (Alexa Flour® 488 or Alexa Flour® 594, Molecular Probes, diluted 1:1000 in block buffer) were incubated with coverslips in a dark, humidified chamber for 1 hour at room temperature. Coverslips were then washed 3 times with wash buffer and stained with 4,6-diaminidino-2-phenylidole (DAPI) at 50ng/ml, mounted in Vectashield (Vector) and sealed with nail varnish. Where cells were treated with Digitonin, media was aspirated, and cells incubated for 10mins on ice in cold PBS. Then cells were incubated in Digitonin buffer (110mM KoAc, 25mM K-HEPES, 2.5mM MgCl_2_, 1mM EGTA) with 0.01% Digitonin for 5mins on ice, washed in room temperature PBS and fixed with 4% paraformaldehyde as above.

### Proximity Ligation Assay (PLA)

PLA assay was performed following Duolink® PLA Fluorescence Protocol (Sigma-Aldrich). Cells were grown, fixed and permeabilised as for immunofluorescence. Coverslips were then incubated for 1 hour with PLA block buffer provided with the Duolink in situ PLA probes at room temperature and incubated with primary antibodies as for immunofluorescence. Coverslips were then washed 3 times with Duolink wash buffer A and Duolink probes incubation, ligation and PLA signal amplification was performed using Duolink® In Situ Detection Reagents Red kit according to the manufacturer’s instructions. In the last wash coverslips were incubated with DAPI at 50ng/ml in 0.01x wash buffer B for 5min, mounted in Vectrashield and sealed with nail varnish.

### FRET-FLIM

HeLa cells transiently expressing GFP-NBAS with mCherry or mCherry-UPF1 were grown on slides for 48 post transfection, fixed in 4% PFA for 10 min, washed in PBS and mounted in Vectashield with DAPI to visualize cell nuclei. Fluorescence lifetime images were acquired on a Leica SP5 SMD confocal laser-scanning microscope fitted with a time-correlated single photon counting module (PicoHarp 300) using a 63/1.4 numeric aperture HCX PL Apo oil immersion objective lens, as previously described (Saleeb et al., 2019). The donor EGFP was excited using a tunable white light supercontinuum laser operating at 488 nm and pulsing at 40 MHz. Emission was detected with an external single-photon avalanche diode (MicroPhoton Devices). Single-pixel fluorescence lifetime analyses were carried out with SymPhoTime version 5.4.4 (PicoQuant).

### FCS and FCCS

FCS measurements were performed by recording fluorescence intensity fluctuations in a very small, approximately ellipsoidal observation volume element (OVE) (about 0.2*μm* wide and 1μ*m* long) that is generated in HeLa cells by focusing the laser light through the microscope objective and by collecting the fluorescence light through the same objective using a pinhole in front of the detector to block out-of-focus light. The fluorescence intensity fluctuations caused by fluorescently labeled molecules passing through the OVE were analyzed using temporal autocorrelation analysis. HeLa cells were mock-depleted or depleted of NBAS in two rounds of depletions as described above. During the second round of depletions cell were co-transfected with 1µg of mCherry control or of mCherry-UPF1plasmids. For NBAS overexpression, cells were co-transfected with GFP-NBAS and mCherry or mCherry-UPF1 (1µg of each). Prior to FCS/FCCS measurement, cells were grown overnight on chambered coverslips (μ-slide, 8 well, Ibidi) and the growth medium was replaced with L-15 medium (Leibovitz) (Sigma-Aldrich L1518) immediately prior to FCS/FCCS measurements. Where appropriate, ER was visualized by the addition of 1µM ER Tracker (green) dye (Thermo Fisher, E34251). Measurements were made only in weakly expressing cells. Fluorescence microscopy imaging of HeLa cells and FCS measurements were performed on a PicoQuant modified Leica SP5 microscope which features Avalanche PhotoDiodes that enable close to single-photon detection (Vukojević et al., 2008). Fluorescence intensity fluctuations were recorded for 50s or 100s. Autocorrelation curves were analysed using the SymphoTime 64 software package (PicoQuant). Control FCS measurements to assess the detection volume were routinely performed prior to data acquisition, using dilute solutions (10nM and 20nM, respectively) of Alexa488 and Alexa568 dyes (Petrášek and Schwille, 2008). The variability between independent measurements reflects the variability between cells, rather than imprecision of FCS measurements. In order to ascertain that the interpretation and fitting of FCS curves is correct, we have: (1) tested several laser intensities in our FCS measurements and have utilized the highest laser intensity, for which the highest counts per second and molecule (CPSM) were obtained, while photobleaching was not observed; and (2) we have established that CPSM do not change among FCS measurements performed. Moreover, we have previously shown that both characteristic decay times increase when the size of the OVE is increased (Fig. 4 in (Vukojević et al., 2008)). Together, these lines of evidence indicate that both short and long characteristic decay times are generated by molecular diffusion rather than by photophysical and/or chemical processes such as protonation/deprotonation of fluorescent proteins. (3) We have ascertained that the long characteristic decay time of our FCS measurements is not the result of photobleaching and that differences in the relative amplitudes of the fast and slow diffusing components reflect differences in their concentrations among cells.

## QUANTIFICATION AND STATISTICAL ANALYSIS

For information about the number of replicates, the meaning of error bars (e.g., standard error of the mean) and other relevant statistical analysis please see the corresponding figure legend. For information about how data was analyzed and/or quantified, please see the relevant section in METHOD DETAILS and/or the figure legend. GraphPad Prism software was used for the statistical analysis in Figures 1C, 1D, 3A, 4B, 5C, 6C, 6E, S1B, S1C, S3A, S3B, RStudio was used for the statistical analysis in Figures 2A, 2B, 2D and S2A. For the specific packages and algorithms used see the ‘‘Software and Algorithms’’ section of the Key Resources Table.

## Data availability

All RNA-Seq data will be available at GEO. Accession number being processed.

The raw uncropped gels, blots and confocal micrographs will be deposited at Mendeley Data.

