## Supplemental Figures for "Identification of a Nonsense-Mediated Decay pathway at the Endoplasmic Reticulum"

Running title: NMD regulation at the ER

[*Keywords:* Nonsense-mediated decay (NMD); RNA quality control; UPF1; NBAS;

Syntaxin 18; ER stress, UPR]

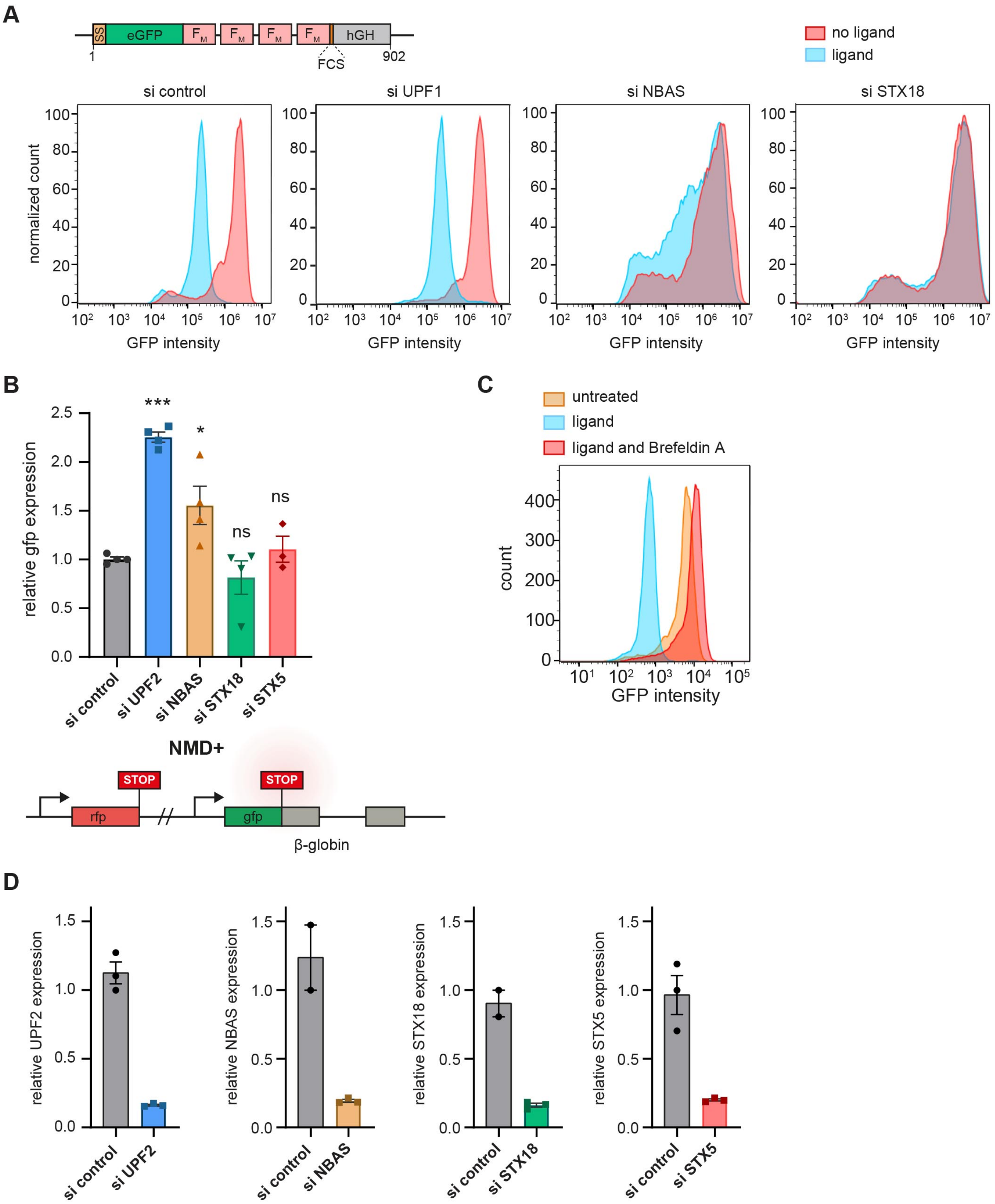
A
SS
eGFP
FM
FM
FM
FM
hGH
1
FCS
902
no ligand
ligand
si control
si UPF1
si NBAS
si STX18
normalized count
GFP intensity
B
relative gfp expression
***
*
ns
ns
si control
si UPF2
si NBAS
si STX18
si STX5
NMD+
STOP
STOP
rfp
gfp
β-globin
C
untreated
ligand
ligand and Brefeldin A
count
GFP intensity
D
relative UPF2 expression
relative NBAS expression
relative STX18 expression
relative STX5 expression
si control
si UPF2
si NBAS
si STX18
si STX5

**Figure S1. Related to Figure 1.**

**Independent functions of NBAS in NMD and Golgi-to-ER transport**.

(A) Top panel shows a schematic representation of the GFP-based reporter expressed in C1 HeLa cells used to measure constitutive secretion. SS: signal sequence; eGFP: enhanced green fluorescent protein; FM: FKBP mutated; FCS: furin cleavage sequence; hGH: human growth hormone, and numbers represent amino acid residues. HeLa C1 cells that constitutively express this reporter are unable to secrete eGFP because the mutant FKBP proteins (FM) form ligand-reversible aggregates that are retained at the ER. The addition of ligand causes aggregate solubilization and rapid eGFP secretion that can be measured by flow cytometry (Gordon et al., 2010). Defects in secretion are manifested by the inability of cells to lose GFP fluorescence even after the addition of ligand. Lower panels show examples of flow-cytometry before (in red), and after the addition of ligand (in blue), for control-depleted cells, cells depleted of NMD factors UPF1 and NBAS, and of secretion factor, STX18.

(B) NMD activity is not affected by the depletion of secretion factors STX18 and STX5. HeLa cells were transfected with a fluorescent NMD+ reporter (gift from K. Lukyanov) and depleted of NMD factors UPF2 or NBAS, or secretion factors STX18 or STX5. Cells carrying the NMD+ reporter were identified by flow cytometry by the presence of red fluorescence, whereas NMD activity was determined by the mean green fluorescence in all red cells. Depletion of both NMD factors increased the mean green fluorescence in comparison to mock-depleted cells, as expected, whereas STX18 and STX5 depletion did not affect the mean green fluorescence. Each point represents one biological replica, bars indicate mean with SEM. Significance was determined by two-tailed unpaired t-test: ***: $p<0.0001$; *: $p<0.05$; ns: not significant.

(C) Brefeldin A treatment blocks constitutive secretion. HeLa C1 cells carrying a GFP-based secretion reporter were FACS-sorted to monitor their ability to secrete GFP. C1 cells (in orange) were able to secrete a GFP-based reporter following addition of ligand (in blue). Brefeldin A treatment led to a block in secretion and a concomitant accumulation of GFP even in the presence of ligand (in red).

(D) Representative efficiency of depletion of NMD and secretion factors was determined by qRT-PCR relative to mock-depleted cells. In each case, relative expression of the depleted factor was normalized to POLR2J. Each point represents one biological replica, bars indicate mean with SEM.

A

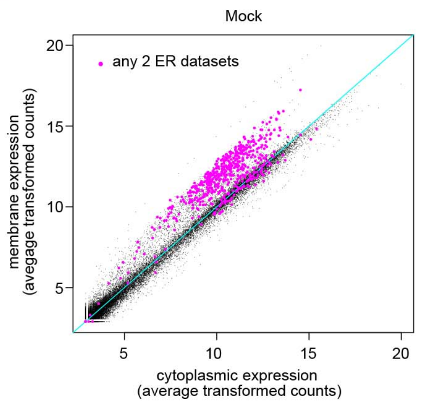


B

| n# membrane- associated genes | 1999 | % regulated |
|---|---|---|
| up in siNBAS mem fraction | 356 | 17.81% |
| up in siUPF1 mem fraction | 788 | 39.42% |
| up in siUPF2 mem fraction | 132 | 6.60% |
| **n# non-membrane genes** | **4194** | |
| up in siNBAS cyto fraction | 147 | 3.51% |
| up in siUPF1 cyto fraction | 1255 | 29.92% |
| up in siUPF2 cyto fraction | 702 | 16.74% |

C

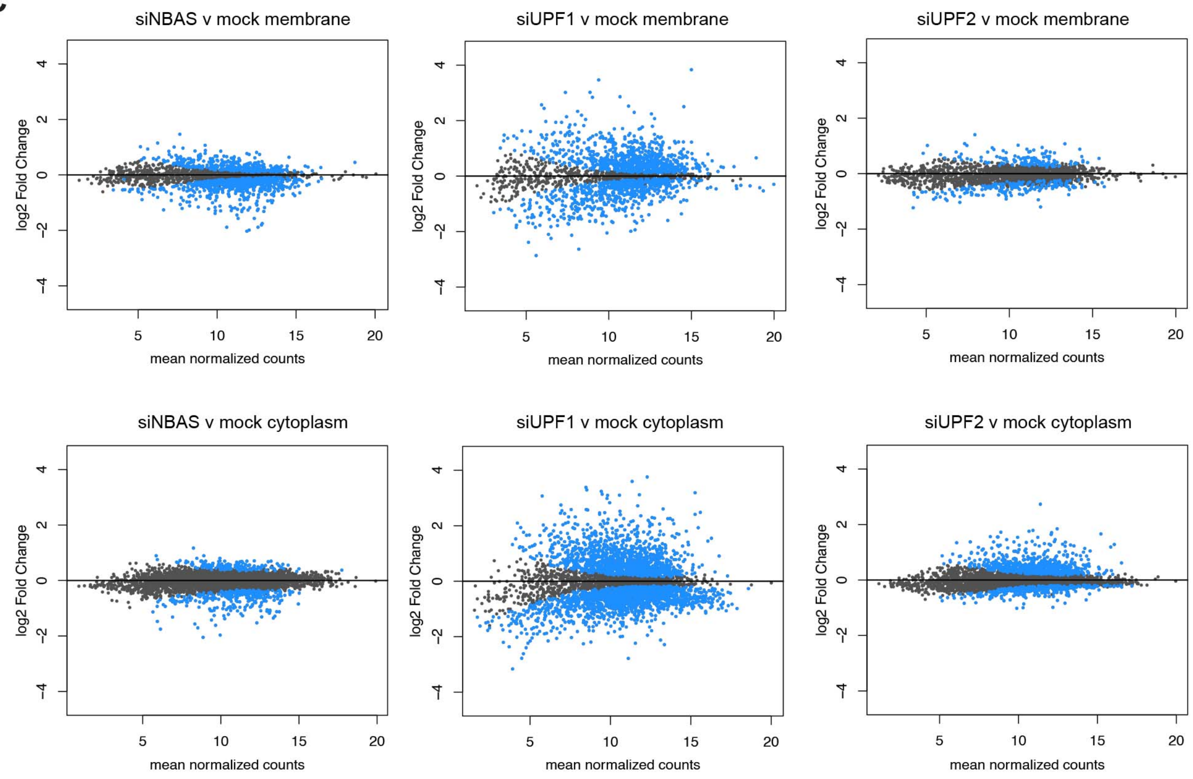


**Figure S2. Related to Figure 2.**

**NBAS regulates membrane-associated RNA targets.**

(A) Experimentally validated ER genes are present in the membrane fraction. Scatter-plot of gene expression in membrane vs cytoplasmic fractions in mock-depleted cells, with experimentally validated ER genes present in 2 out of 3 datasets described in panel 2B are indicated in magenta. (OR = 66.94, p-value < 2.2e-16, Fishers Exact test)

(B) Summary table of genes regulated by NMD factors in each cellular fraction. Genes were defined as membrane-enriched or non-membrane-enriched as described in figure 2C. Genes were classified as regulated if they were significantly ($p<0.05$) upregulated in that fraction when the NMD factor was depleted.

(C) Differential expression analysis of genes regulated by NMD factors in membrane fractions and in the cytoplasm. MA plots show log2 fold-change of gene expression plotted over mean of normalized counts (gene expression). Significantly changed genes ($p<0.05$) are indicated in blue.

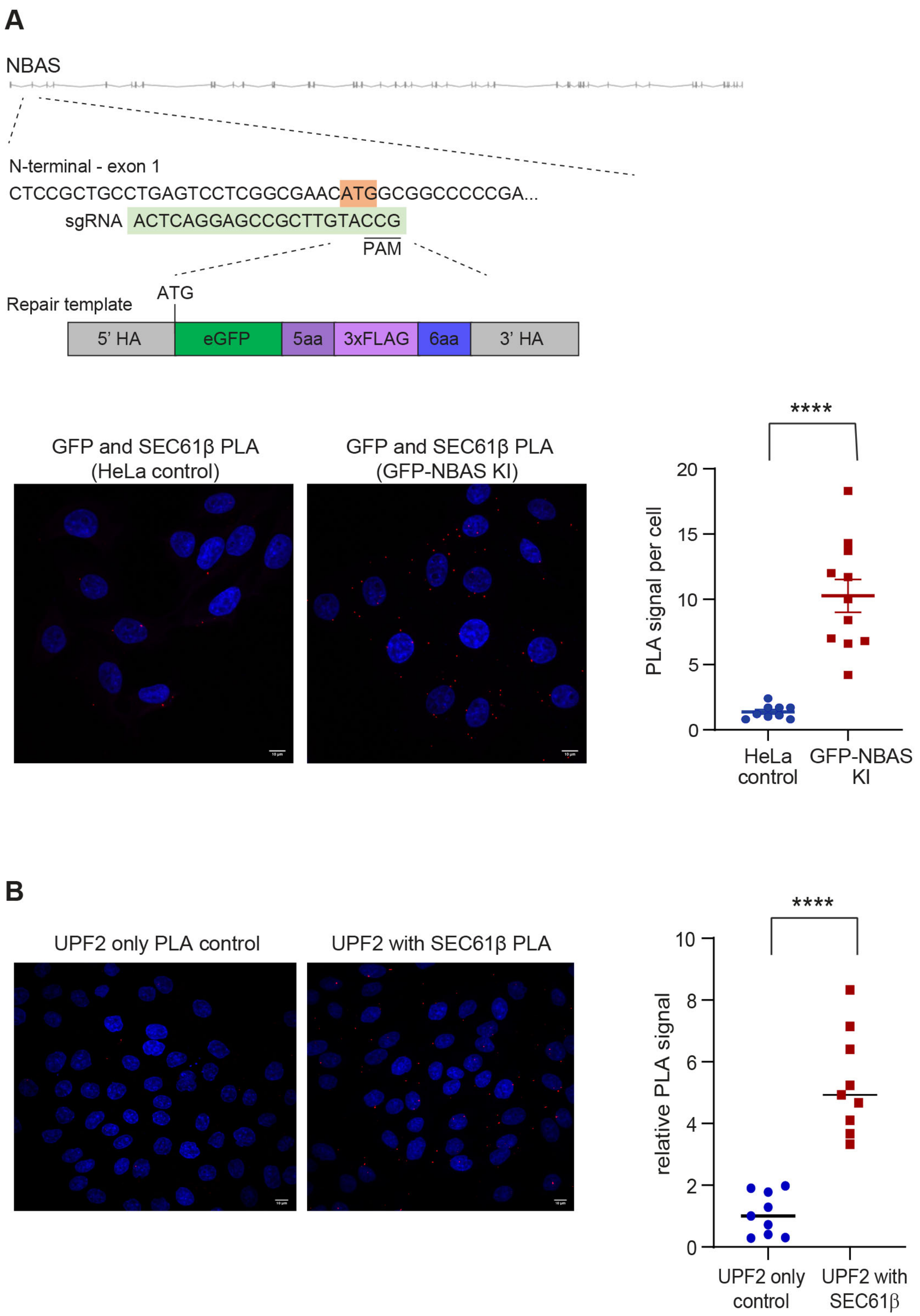


**Figure S3. Related to Figure 4.**

**Localization of endogenous NBAS and UPF2 proteins to the ER.**

(A) Top panel depicts the strategy used to knock-in an eGFP/3xFLAG tag at the N-terminus of the NBAS locus, using CRISPR/Cas9-mediated genome editing in HeLa cells. Direct interaction between endogenous epitope-tagged NBAS and SEC61β proteins was determined by PLA using anti-GFP and anti-SEC61β antibodies, in HeLa KI (epitope tag knock-in) cells or wild-type HeLa cells, as a control. The graph shows the quantification of PLA signal. Each point represents mean PLA count per cell in one captured frame, relative to the HeLa negative control. Significance was determined by Mann-Whitney test: ****: $p<0.0001$.

(B) UPF2 is localized in the close proximity of the SEC61 translocon component at the ER. Direct interaction between endogenous UPF2 and SEC61β proteins was determined by PLA using the anti-UPF2 and anti-SEC61β antibodies, in HeLa cells. The graph shows the quantification of PLA signal. Each point represents mean PLA signal per cell in one captured frame, relative to the HeLa negative control. Significance was determined by Mann-Whitney test: ****: $p<0.0001$.

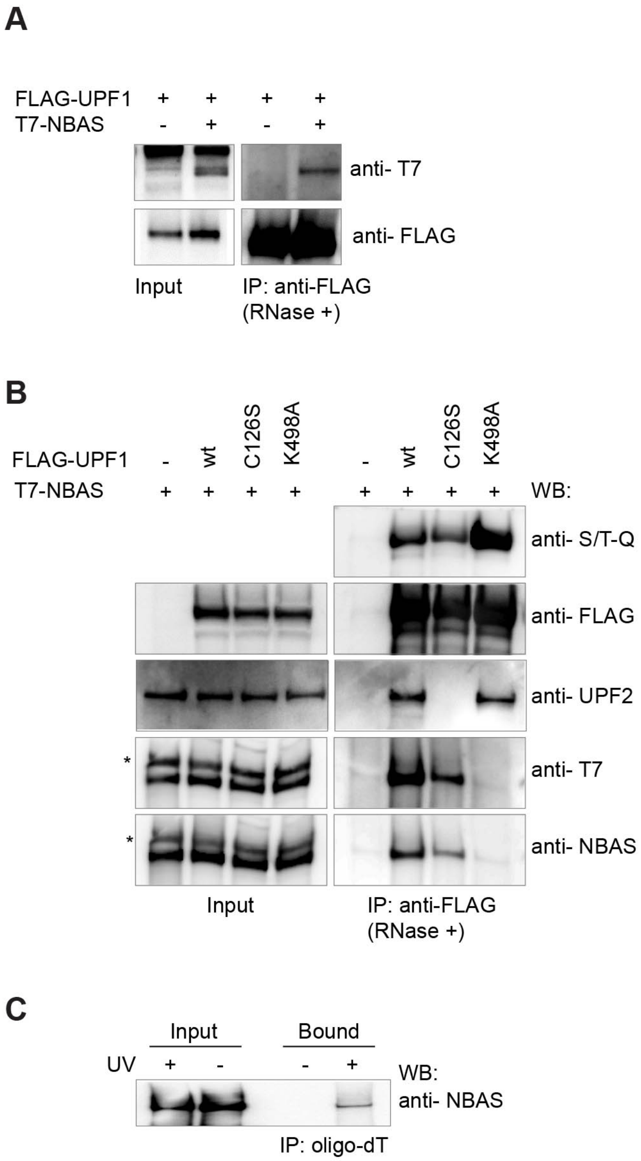


**Figure S4. Related to Figure 5**

**NBAS interacts with UPF1 and preferentially associates with the SURF complex.**

(A) T7-tagged NBAS was co-expressed with FLAG-tagged UPF1 in HeLa cells. Cell lysates were subjected to immunoprecipitation with anti-FLAG antibody in the presence of RNase. The NBAS-UPF1 interaction was revealed by Western blot with anti-T7 antibody.

(B) T7-NBAS was co-expressed in HeLa cells with FLAG-tagged wt UPF1; C126S hypo-phosphorylated mutant UPF1 predominantly present in the SURF complex, or with K498A ATPbinding mutant UPF1, which is mainly associated with the DECID complex. Anti-FLAG IPs were performed in the presence of RNase and subjected to Western blot analysis with the indicated antibodies. To detect phosphorylated UPF1, anti-FLAG-IPs were probed with a phospho-(Ser/Thr) ATM/ATR substrate antibody (anti-S/T-Q).

(C) NBAS is bound to mRNAs. Hela cells were UV crosslinked and mRNP complexes were purified using oligo dT beads. The presence of NBAS in mRNP complexes was revealed by Western blot with anti-NBAS antibody.

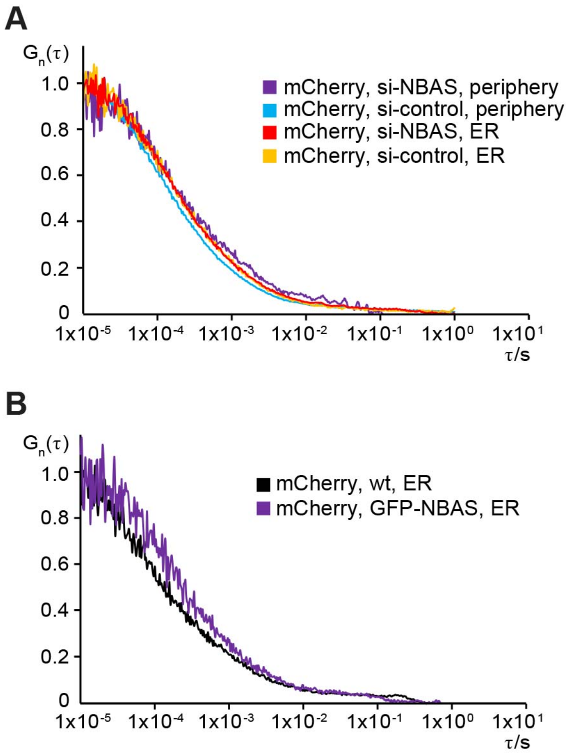


**Figure S5. Related to Figure 6.**

**FCS control experiments**

(A) Average autocorrelation curves obtained for FCS measurements indicate that mCherry alone shows an almost complete overlap of decay times upon depletion of NBAS by RNAi, or between the ER and the cytoplasmic periphery.

(B) Overexpression of GFP-NBAS does not affect the mobility of mCherry control at the ER.
